# A hyper-quiescent chromatin state formed during aging is reversed by regeneration

**DOI:** 10.1101/2023.02.14.528512

**Authors:** Na Yang, James R. Occean, Daniël P. Melters, Changyou Shi, Lin Wang, Stephanie Stransky, Maire E. Doyle, Chang-Yi Cui, Michael Delannoy, Jinshui Fan, Eliza Slama, Josephine M. Egan, Supriyo De, Steven C. Cunningham, Rafael de Cabo, Simone Sidoli, Yamini Dalal, Payel Sen

**Affiliations:** Laboratory of Genetics and Genomics, National Institute on Aging, NIH; Baltimore, MD; Laboratory of Receptor Biology and Gene Expression, National Cancer Institute, NIH; Bethesda, MD; Department of Biochemistry, Albert Einstein School of Medicine; Bronx, NY; Laboratory of Clinical Investigation, National Institute on Aging, NIH; Baltimore, MD; JHU SOM Microscope Facility, Johns Hopkins University; Baltimore, MD; Computational Biology and Genomics Core, Laboratory of Genetics and Genomics, National Institute on Aging, NIH; Baltimore, MD; Department of Surgery, Ascension Saint Agnes Hospital; Baltimore, MD; Translational Gerontology Branch, National Institute on Aging, NIH; Baltimore, MD

**Keywords:** Chromatin, epigenetics, aging, liver, regeneration

## Abstract

Epigenetic alterations are a key hallmark of aging but have been limitedly explored in tissues. Here, using naturally aged murine liver as a model and extending to other quiescent tissues, we find that aging is driven by temporal chromatin alterations that promote a refractory cellular state and compromise cellular identity. Using an integrated multi-omics approach, and the first direct visualization of aged chromatin we find that globally, old cells show H3K27me3-driven broad heterochromatinization and transcription suppression. At the local level, site-specific loss of H3K27me3 over promoters of genes encoding developmental transcription factors leads to expression of otherwise non-hepatocyte markers. Interestingly, liver regeneration reverses H3K27me3 patterns and rejuvenates multiple molecular and physiological aspects of the aged liver.

## Introduction

The epigenome is responsible for connecting the genetic information in our chromosomes to functional outcomes; that is, it links the genotype with the phenotype (Berger et al., 2009). During senescence and aging, marked alterations occur in the epigenome including changes in chromatin accessibility, DNA methylation, histone occupancy, histone modification, and 3-dimensional (3D) genome topology (Benayoun et al., 2015; Pal and Tyler, 2016; Sen et al., 2016; Yang and Sen, 2018). Age-related epigenomic changes drive organ dysfunction through increased inflammation, fibrosis, reduced regeneration, and incidence of disease. Given the reversible nature of epigenomic changes, they are attractive therapeutic targets for age-associated decline and disease.

Aside from DNA methylation, age-related chromatin mechanisms have been investigated limitedly, assayed primarily in model organisms such as yeast, worms, flies, or cellular models such as senescent cells in culture, stem cells, or peripheral blood cells (Sen et al., 2016). Consequently, this has restricted our understanding of cellular aging in quiescent mammalian tissues. Quiescence refers to a reversible G0 state of differentiated cells in a tissue required to achieve and maintain optimal organ mass, geometry, and homeostatic function. Quiescent cells do not allocate resources to cell division or growth but remain metabolically active and responsive to various environmental stresses (Fiore et al., 2018). Aged tissue might represent a system where cells have remained quiescent for prolonged periods. However, whether quiescence maintenance in the long-term (a state we call hyper-quiescence in this study) incurs temporal alterations in the epigenomic landscape and impacts organ function is unknown. Alternatively, aged tissues also accumulate senescent cells (Ogrodnik et al., 2017); senescence represents a presumed irreversible non-dividing state distinct from classic quiescence. Of note, senescent cells in most tissues are rare and organ dysfunction is evident even before senescent cells can be detected, suggesting that prolonged quiescence may play a dominant role in age-associated functional decline.

One chromatin mechanism that has been implicated but not investigated deeply in quiescence and aging is lysine trimethylation on histone H3 (H3K27me3; reviewed in (Kane and Sinclair, 2019)). H3K27me3 is a key chemical modification on chromatin that induces compaction and gene silencing. This repressive histone mark is “written” by polycomb repressive complex 2 (PRC2), which assembles with one of two paralogous catalytic subunits: enhancer of zeste homolog 1 (EZH1) or EZH2 to form canonical and non-canonical complexes (Di Croce and Helin, 2013; Margueron et al., 2008; Yu et al., 2019). PRC2 activity represses key developmental gene promoters required to establish a specific cell lineage. Unlike most repressors, PRC2 is not recruited to these sites by any transcription factor (TF) but rather by the unique chromatin environment. Once these repressive domains are established in development, PRC2 performs heavy maintenance functions (Yu et al., 2019). Interestingly, transcription itself can evict PRC2, suggesting that transcriptionally dormant states (such as prolonged quiescence) can perpetuate modifications such as H3K27me3 (Riising et al., 2014; Yu et al., 2019).

We use the liver as a model organ to investigate chromatin changes in aging given its pivotal role in metabolism and evidence of clear age-related structural and functional decline. Furthermore, the liver has remarkable regenerative capacity (Forbes and Newsome, 2016), which provides an opportune rejuvenation system to test the reversibility of age-associated chromatin states. Our findings uncover H3K27me3 as a central regulator of the aging process and further suggest that regeneration mitigates age-related H3K27me3 patterns. Overall, this work ideates a new concept of cell proliferation-mediated resetting of the epigenomic landscape to reverse aging. Analogous chromatin changes occur in other organs (kidney, heart, and muscle).

## Results

### Increased H3K27me3 is a signature of aging

We used liquid chromatography tandem mass spectrometry (LC-MS/MS) to unbiasedly survey the altered epigenome in the aging liver (n=3 biological replicates per group, Table S1-S2, MS raw data at chorusproject.org/1769). As proof-of-principle, the ratio of variant H3.3 to canonical H3.1/3.2 (hereon H3) was increased in old livers (Fig. S1A), consistent with previous reports (Tvardovskiy et al., 2017). There was no significant difference in the overall abundance of monomethyl (me1), dimethyl (me2), trimethyl (me3) or acetyl (ac) -ated histones with age (Fig. S1B). However, quantification of single histone post-translational modifications (hPTMs) revealed some striking changes with age.

In pairwise comparisons of young and old livers, the repressive hPTMs H3K27me3, H3K18me1, H4K20me2 and a single active mark, H3K18ac, were significantly enriched in old livers (Fig. 1A, red circles). Notably, although H3K18ac was significantly enriched, it showed only a modest fold change. We were intrigued by the increase of H3K27me3 in aging, which has been noted (but not always emphasized) in several previous reports including those in aged *Drosophila* head and muscle (Ma et al., 2018), killifish brain (Baumgart et al., 2014), killifish muscle (Cencioni et al., 2019), aged murine muscle stem cells (Liu et al., 2013; Schworer et al., 2016) and aged human post-mortem brains (Nativio et al., 2020). As examples, we mined the published LC-MS/MS data from murine muscle stem cells and human brains. Much like the aged murine liver, these tissues showed significant increase in H3K27me3 with age (Figs. S1C-D). We used multiple orthogonal methods such as western blot (Fig. 1B), immunofluorescence (IF, Fig. S1E) and Chromatin Immunoprecipitation coupled to sequencing (ChIP-seq) (Fig. S1F, explained in the next section) to validate the increase of H3K27me3 in mouse livers. Furthermore, we carefully examined the hepatocyte ultrastructure by transmission electron microscopy (TEM) coupled to immunogold labeling to visualize the spatial location of H3K27me3 in young and old livers (n=2 biological replicates per group, Table S1). Sections with no primary antibody were used as control to verify the specificity of antibody labeling and indeed most gold particles were detected in the nucleus (Fig. 1C-D). In agreement with LC-MS/MS, western blot, IF and ChIP-seq data, we observed significantly higher gold particles in the nucleus of old hepatocytes (Fig. 1D). Within the nucleus, there was a clear preference for localization in the nuclear matrix (>200 nm from the nuclear membrane) over nuclear periphery (<200 nm of the nuclear membrane, Fig. 1E). Interestingly, the immunogold particles in the old hepatocyte nuclei formed clusters (i.e., 3 or more gold particles in proximity, Figs. 1C and 1F) suggesting an interaction between H3K27me3-rich regions in the aged cell. Together, our results show that a global increase in H3K27me3 is a common epigenetic signature of aging.

**Fig. 1:**
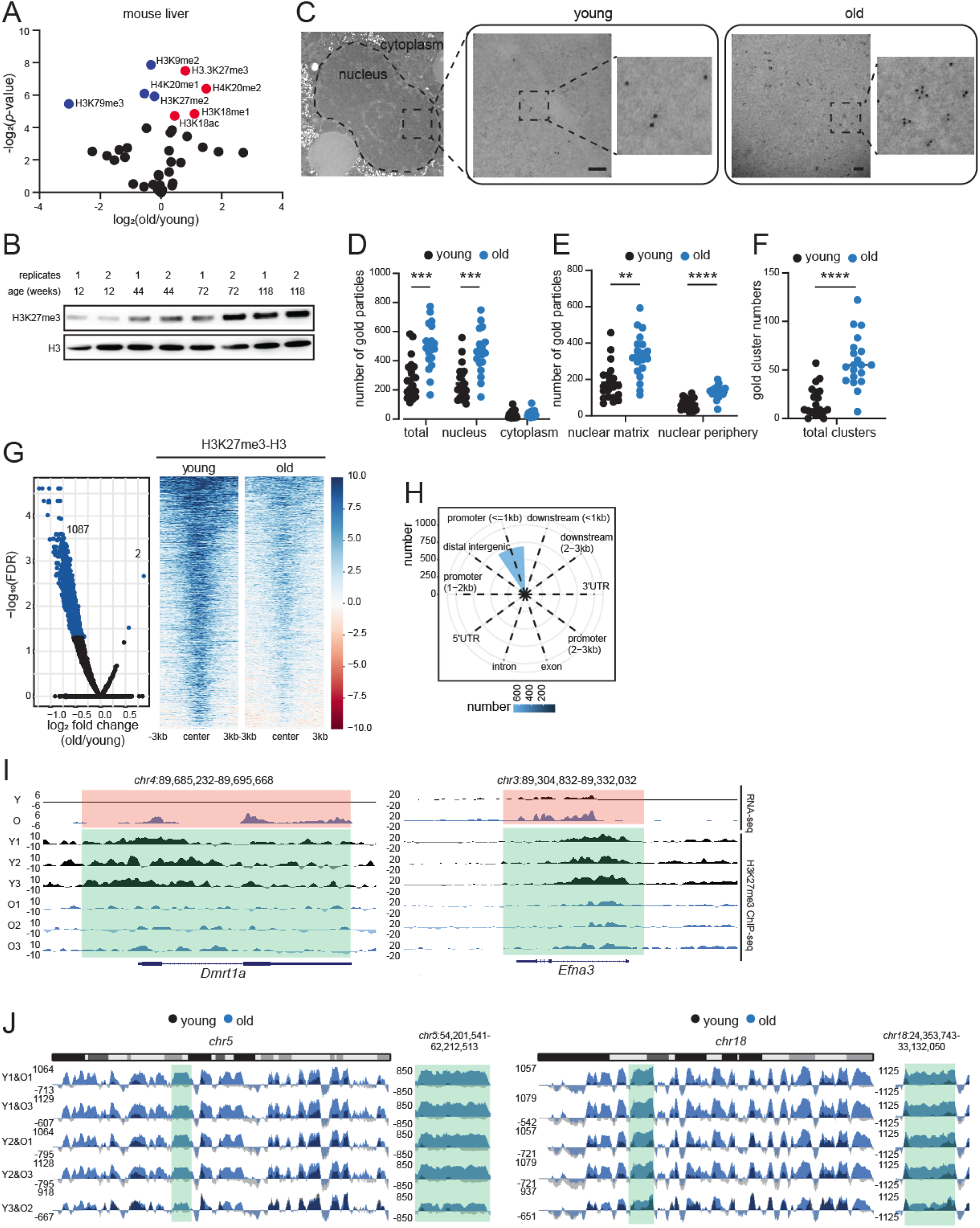
H3K27me3 increases during aging and forms age-domains. **(A)** Volcano plot of single hPTMs in old vs young livers. The significantly increased (red) and decreased (blue) modifications are labeled (two-tailed unpaired t-test). **(B)** Western blot of H3K27me3 and H3 in liver lysates from mice of different ages as indicated (n=2 biological replicates per age group). **(C)** Representative TEM images of young and old hepatocytes with immunogold labeling of H3K27me3 (n=2 biological replicates per group). Scale bar is 100 nm. **(D)** Subcellular location of gold particles. **(E)** Subnuclear location of gold particles. **(F)** Quantification of clusters (i.e., ≥3) of gold particles. Data are summarized as mean ± S.E.M with each dot representing one cell. ** p<0.01, *** p<0.001 and **** p<0.0001 from a Kolmogorov-Smirnov test with corrections for multiple comparisons. **(G)** Volcano plot (left) and heatmap (right) of DiffBind output. Significantly different H3K27me3 peaks (FDR<0.05) are indicated in blue. **(H)** Annotation of differential peaks identified in (G). **(I)** Genome browser snapshots of two differential peak regions from (G). Green regions show age-related loss of H3K27me3, and orange regions show de-repression of genes. **(J)** Genome browser shot of overlayed H3K27me3 ChIP signal (H3 subtracted and E. coli spike-in normalized) over chr5 (left) and chr18 (right) in sex-matched pairs of young and old mouse livers. Green area is expanded on the right of each chromosome.

### H3K27me3 shows a unique genomic pattern in aged tissue: loss of peaks and gain of domains (age-domains)

We next investigated the genomic localization of H3K27me3 in aged livers. We performed H3K27me3 ChIP-seq in young and old livers with total H3 and IgG as controls (n=3 biological replicates per group, Table S1, QC metrics in Fig. S2A-G). Two % exogenous human chromatin from HeLa cells was spiked in before immunoprecipitation for quantitative comparisons (ChIP-Rx (Orlando et al., 2014)). A lower % of human reads were recovered in old H3K27me3 ChIP samples (1.7% in old vs 2.1% in young) suggesting proportionally higher mouse H3K27me3 in old livers (Fig. S1F). In contrast, the recovery of human reads in H3 and input was comparable between the two groups.

A Principal Component Analysis (PCA) of both the overall genome coverage (Fig. S1G) and all H3K27me3 peaks (Fig. S1H), clearly segregated young and old samples suggesting distinct genome enrichment patterns. Interestingly, there were fewer H3K27me3 peaks in the old samples (mean in young 5211 vs old 3354) although the differences were not significant (Fig. S1I). Curiously, the peaks identified in old samples were generally broader, had higher tag density and lower peak height (Fig. S1J) suggesting a “spreading” of the signal. Differential peak analysis revealed only 2 peaks with higher H3K27me3 enrichment and 1087 peaks with lower H3K27me3 enrichment in old livers (FDR<0.05, Fig. 1G blue dots and heatmap, Table S3). These differentially enriched peaks were annotated primarily to promoters (Fig. 1H). Example genome browser views of two such annotated regions and related gene expression change are shown in Fig. 1I. These findings however were inconsistent with our LC-MS/MS, IF, TEM, western blot and ChIP-seq spike-in results that showed a global accumulation of H3K27me3 with age.

The global increase in H3K27me3 (Fig. 1), the clear separation of H3K27me3 genome coverage (Fig. S1G), and the spreading of local H3K27me3 signal (Fig. S1J) in old livers prompted us to look deeper into H3K27me3 genome distribution patterns to address the inconsistency. The full extent of H3K27me3 signal was captured in the whole chromosome view. At this scale, old livers showed a strong and reproducible enrichment across multi-megabase sized regions across all chromosomes (chr5 and chr18 shown as examples in Figs. S3-S4). We call this type of enrichment “age-domains”. Age-domains were found in all 3 old biological samples (Figs. S3C and S4C). The enrichment was more prominent when we scaled the coverage maps to spike-in reads (Fig. 1J). In sum, our ChIP-seq data identified two different chromatin states based on H3K27me3 genome-enrichment patterns: young (more peaks, less age-domains) and old (less peaks, more age-domains).

### Age-domains are heterochromatinized while H3K27me3 peak regions are euchromatinized during aging

To assess whether the H3K27me3 peaks and age-domains segregate into different fractions of chromatin, we measured the salt solubility of mononucleosomes from these regions by extraction under different salt concentrations (Fig. 2A). The extracted DNA was purified and analyzed by sequencing. The final pellet (insoluble chromatin) was dissolved in a solubilization buffer and analyzed in a similar fashion (n=2 replicates per group, Table S1, QC metrics in Fig. S5A-G). As has been reported previously (Henikoff et al., 2009), we noted that low salt concentrations (0-250 mM) extracted very few histones in the supernatant (Fig. 4F) and the associated DNA mapped to active regions of the genome (Fig. 2B inset 1 and Fig. S6 inset 1). In the young samples, the euchromatin eluted in the 150 and 250 mM fractions. Euchromatic regions in the old liver showed relatively lower enrichment (Fig. 2B inset 1 and Fig. S6 inset 1) suggesting a decrease in its proportion with aging. Importantly, the old euchromatin fraction eluted at lower salt concentrations (0, 67.5, and 150 mM salt) suggesting a decondensation of this fraction with aging. H3K27me3 peaks in young that were lost in the old (Fig. 1G) were among those that showed chromatin decondensation (Fig. 2C). Remarkably, we observed that H3K27me3 age-domains in old livers overlapped almost perfectly with the 350 mM and pellet fraction (Fig. 2B inset 2, Fig. S6 inset 2 and Fig. 2D) suggesting that these regions are packed into insoluble heterochromatin.

**Fig. 2:**
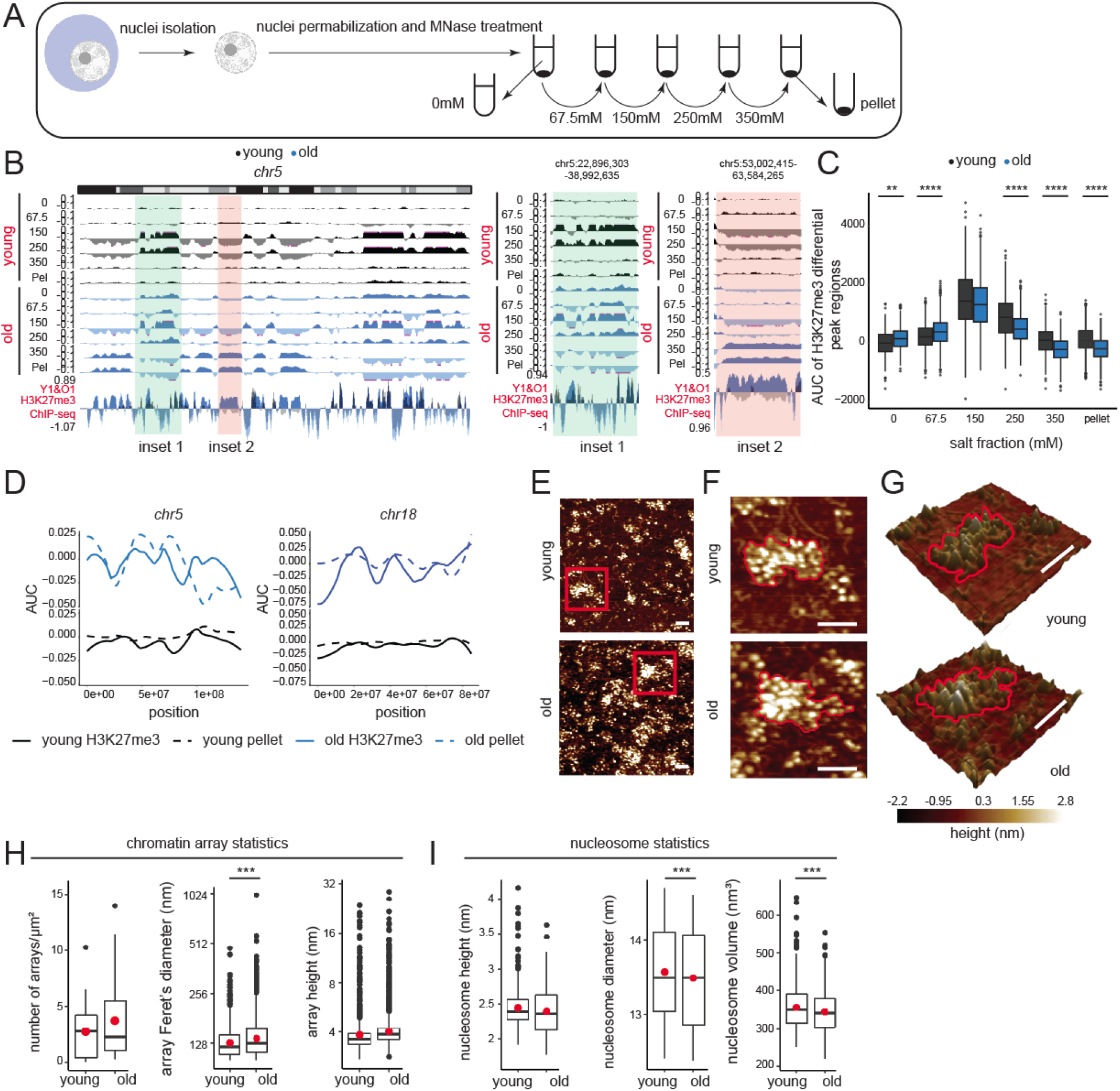
Age-domains are heterochromatinized while H3K27me3 peak regions are euchromatinized during aging. **(A)** Schematic of salt fractionation experiment (n=2 replicates per group). **(B)** Genome browser snapshot of salt fraction enrichments on chr5. Inset 1 (green) is euchromatinized and inset 2 (orange) is heterochromatinized with age and expanded on the right. **(C)** The salt elution profile of differential peaks in Fig. 1G. ** p<0.01, **** p<0.0001 from a Welch’s t-test. **(D)** H3K27me3 AUC (solid line) and salt fraction (dotted line) over chr5 and chr18 depicted as fraction of maximum. **(E)** Representative AFM images of chromatin arrays extracted with 67.5 mM salt (n=3 biological replicates per group). **(F)** The red boxed region in (E) is magnified. An array in each sample is highlighted with a red contour. **(G)** Same as (F) but represented as a 3D image. For (E-G), scale bar is 100 nm. **(H)** Chromatin array number per µm^2^, Feret’s diameter, and array height. **(I)** Nucleosomal height, diameter, and volume. For H-I, the red dot represents the mean. *** p<0.001 from a Welch’s t-test.

To physically assess whether differences exist between chromatin structures in young and old livers, we purified chromatin using a very mild micrococcal nuclease (MNase) digestion protocol followed by extraction from nuclei in buffer containing 67.5 mM NaCl. We analyzed these samples (n=3 biological replicates per group, Table S1) by single-molecule atomic force microscopy (AFM). By very mildly digesting chromatin, only the most accessible (hyper-sensitive) DNA is cut, thereby, preserving the more compacted chromatin arrays. This allowed us to quantify how many distinct chromatin arrays were isolated from young and old liver nuclei under identical conditions (Fig. 2E-G). Qualitatively, chromatin obtained from young livers possessed distinct well-organized clusters. In contrast, chromatin from old livers appeared less coherent, with an overall increase in nucleosome number per field and a “carpet-like” appearance (Fig. 2E). This result agrees with the observed increase in solubility of old chromatin at 67.5 mM salt (Fig. 2B-C). Next, we quantified the number of distinct chromatin arrays (clusters) isolated, as well as the height and Feret’s diameter of these arrays. Despite a trend of a larger mean for number and height of arrays in the old livers, these results were not statistically significant (Fig. 2H, panel 1 and 3). However, in chromatin obtained from old livers, the Feret’s diameter was broader (Fig. 2H, panel 2), showing that arrays from old liver nuclei are larger. To delve deeper into nucleosome-level differences, we next quantified the dimensions of individual nucleosomes (height, diameter, and volume) in arrays obtained from young and old livers. Although the height of nucleosomes did not differ (Fig. 2I, panel 1), compared to nucleosomes obtained from young livers, nucleosomes from older livers had smaller diameters, and correspondingly, a smaller volume (Fig. 2I, panels 2 and 3). Taken together, these data suggest that in older samples, individual nucleosomes are more compacted, possibly reflecting a less accessible state of site exposure and that chromatin arrays from older livers occupy larger areas, suggesting either an increased number of nucleosomes per array and/or a loss in higher order coherence.

### Consequences of H3K27me3 genomic redistribution in aging

We wanted to further probe the functional consequences of H3K27me3 peak loss and age-domain gain in old livers. Interestingly, genes near H3K27me3 peaks that were lost in the old livers were related to development (particularly nervous system and cardiac), and cell differentiation (Fig. 3A). Loss of H3K27me3 de-repressed these genes (Fig. 3B). Ontology terms associated with neuronal function were also seen in the genes upregulated with age (Fig. S9D). Most of these genes that lost H3K27me3 signal with age encode for developmental TFs such as *Hox, Ascl1, Sox2, Neurog1, Gata4* etc. (Fig. 3C, Table S3) that are known to play important roles in lineage specification. Indeed, expression of only 3 TFs, one of which is *Ascl1*, can reprogram hepatocytes to neurons (Marro et al., 2011).

**Fig. 3:**
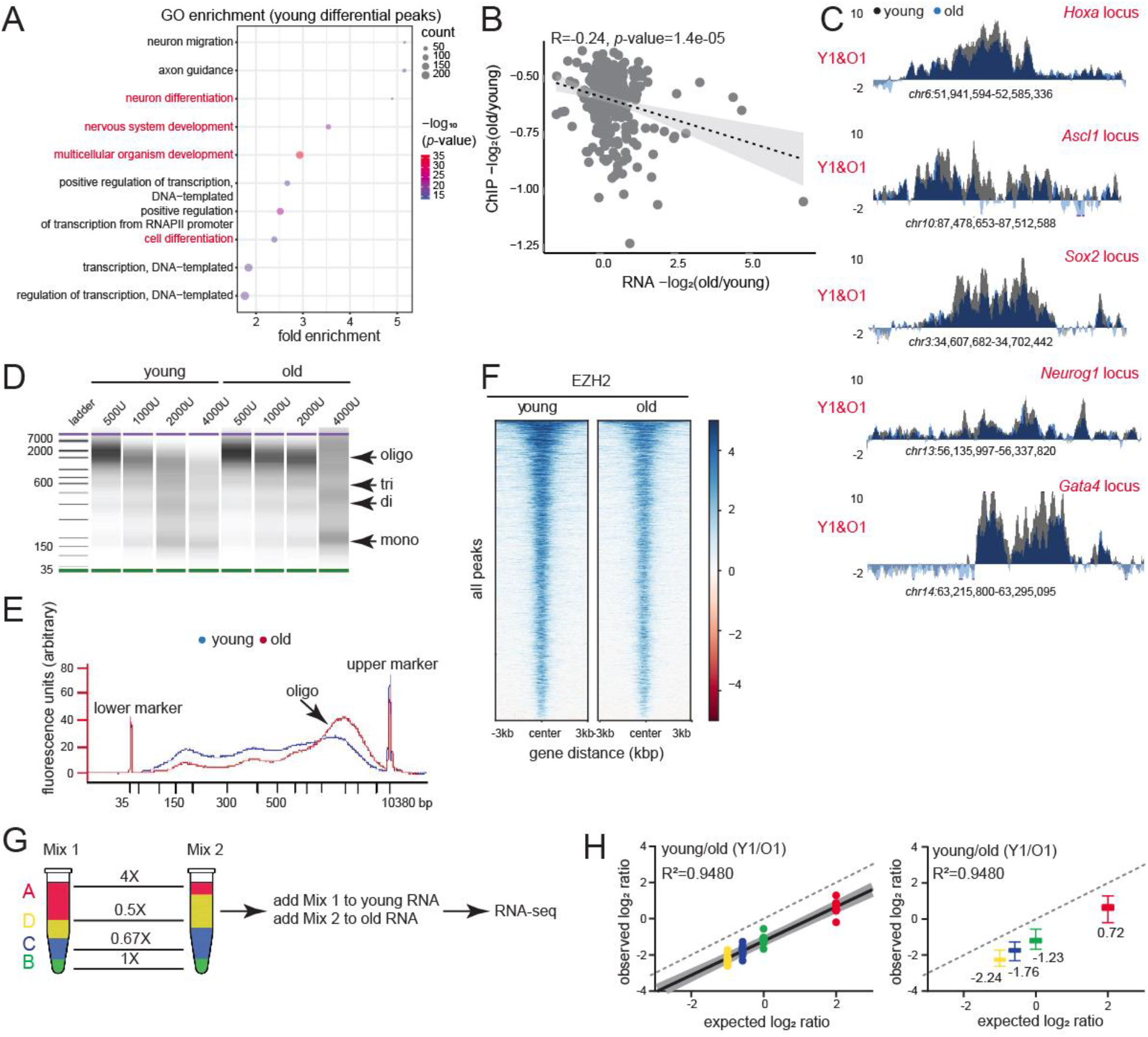
Consequences of H3K27me3 genomic redistribution in aging. **(A)** GO terms associated with differential peaks in Fig. 1G. Development and differentiation terms are indicated in red. **(B)** Negative correlation between H3K27me3 and gene expression change in old vs young. **(C)** H3K27me3 ChIP-seq profiles at indicated loci in young and old livers. **(D)** BioAnalyzer profiles of fixed chromatin from young and old livers digested with increasing units of MNase. **(E)** The digestion profile with 2000U of MNase shown as an overlayed trace. **(F)** Heatmap showing EZH2 signal at all called peaks in young and old livers. **(G)** ERCC Exfold transcript abundance and overall spike-in strategy. A-D represent the 4 groups of transcripts present in Mix 1 and 2. **(H)** Observed vs expected plot of Mix1/Mix2 log ratio for one pair of sex-matched young and old animals before resection. The dotted line represents a hypothetical experiment where the observed Mix1/Mix2 ratio is the same as expected. On the right, same data shown as box plots with median values indicated.

To assess whether H3K27me3-mediated regulation of developmental genes was conserved in aging human livers, we performed Cleavage under targets and tagmentation (CUT&Tag) (Henikoff et al., 2020) with H3K27me3 antibody on human liver core-needle biopsies (n=3 and 5 biological replicates for young and old respectively, Table S1, QC metrics in Fig. S7A-G). These human subjects were hospitalized for either cholecystectomy, bariatric procedures, or gastroesophageal reflux disease, and had otherwise normal livers. Given the limited sample amount, we were unable to perform ChIP-seq. Nevertheless, from CUT&Tag data, we identified age-related differential sites in the young (1117 regions) and old (62 regions) samples (Fig. S7H), which was numerically similar to the changes seen in mice. As in mice, genes annotated to the 1117 regions with lower H3K27me3 signal in the old were linked to neuronal and cardiac development (Fig. S7I). Of note, we could not visualize age-domains of H3K27me3 in our human samples due to the limitation of CUT&Tag in capturing domain-like enrichments. However, our peak level analyses in both mice and humans suggest a major loss of H3K27me3-mediated suppression of cell identity genes.

While local loss of H3K27me3 compromises cellular identity, we surmised that broad heterochromatinization (as seen in salt fractionation and AFM studies) may have more global effects such as reduction of chromatin accessibility and global transcription suppression. In concordance, we noted that old chromatin was in general more resistant to MNase digestion compared to young suggesting the formation of a more condensed structure with age (Fig. 3D-E). This in turn, likely reduces the genome-wide binding of EZH2 to its target regions (Fig. 3F). To ascertain whether the global transcriptome was altered during aging, we prepared RNA-seq libraries with Exfold External RNA Control Consortium (ERCC) spike-in controls (Loven et al., 2012). ERCC spike-ins allow for comparisons across samples with different global RNA levels. We added Mix 1 to young samples and Mix 2 to old samples prior to ribo-depletion and library generation (Fig. 3G). The data were normalized to reads per kilobase per million mapped reads (RPKM) and filtered to remove transcripts <1 RPKM. The log_2_RPKM (Mix1/Mix2) for each ERCC transcript (observed ratio) and the log2 ratio of the known attomolar concentrations of the two mixes (expected ratio) were plotted (Fig. 3H). If global transcription is unaffected, the observed ratio (black solid line with gray shaded 95% confidence bands) of Mix1/Mix2 ERCC transcripts is predicted to mirror the expected ratios (dotted line). Our analysis, however, showed that Mix1/Mix2 ratios were lower than expected, suggesting a strong genome-wide transcription suppression in old livers.

### H3K27me3 patterns in aging are mimicked by deep quiescence cultures (hyper-quiescence)

While EZH2 plays a dominant role in H3K27me3-mediated gene repression in actively dividing cells, the paralogous EZH1 is highly expressed in post-mitotic tissue (Margueron et al., 2008). We hypothesized that as tissues spend more time in the post-mitotic stage, i.e., with age, they switch to a more EZH1-dependent mode of repression. To test this, we measured the levels of enzymes affecting H3K27me3 in young and old livers. At the RNA level there was no age-associated change in expression of EZH1/2 or demethylases KDM6A/6B (Fig. 4A, Table S1, n=8 biological replicates per group). However, at the protein level, EZH2 levels declined while EZH1 showed increase with age (Fig. 4B). Protein levels of the demethylases KDM6A/6B remained unaltered with age (Fig. 4B). Of note, KDM6A is encoded by the X chromosome and hence is about half the level in males compared to females. KDM6C was not tested as it is a male-specific inactive paralog. Our results support an EZH2-to-EZH1 switch mechanism in aging.

**Fig. 4:**
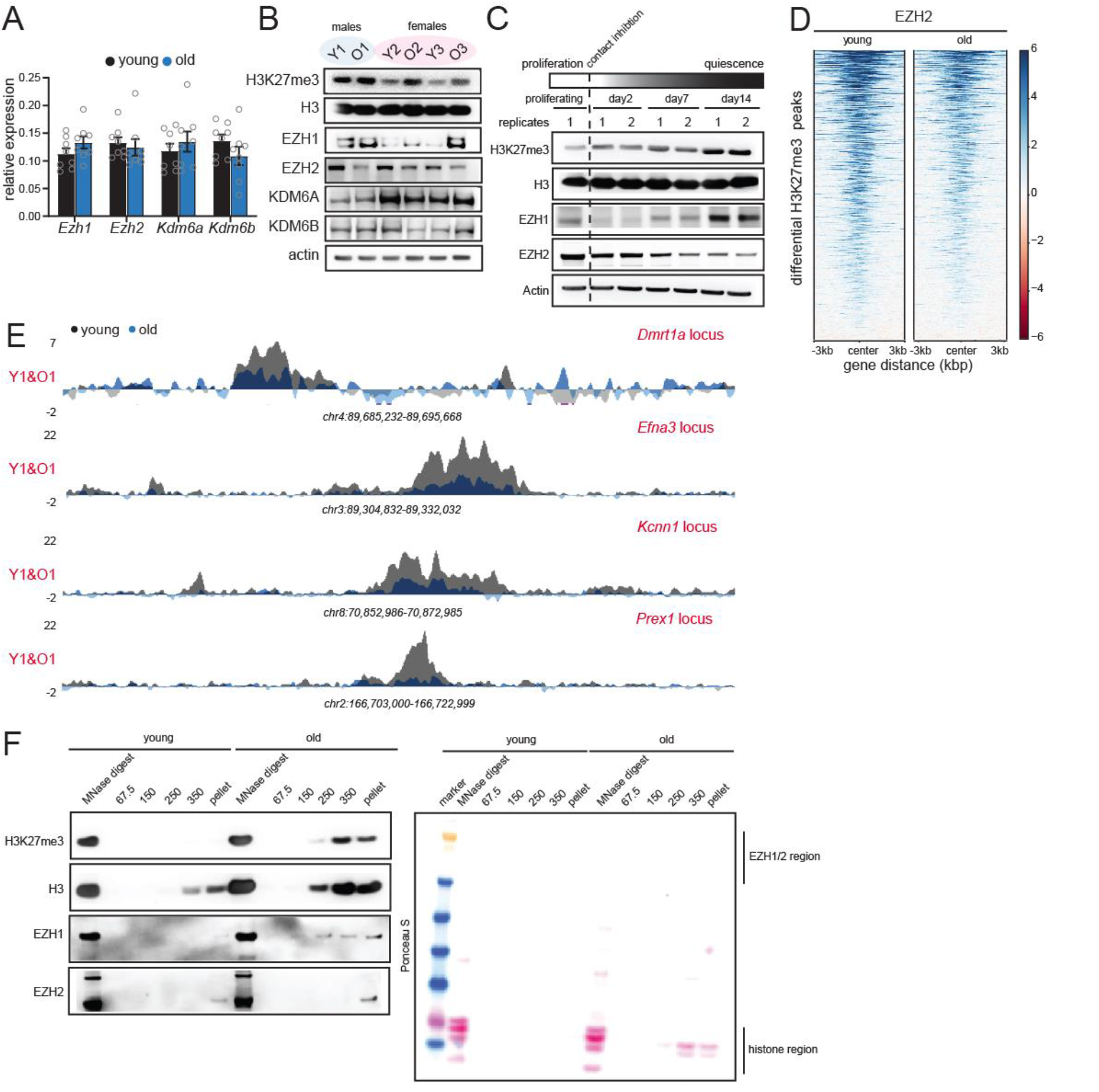
H3K27me3 patterns in aging are mimicked by deep quiescence cultures (hyper-quiescence) **(A)** RT-qPCR analysis of *Ezh1, Ezh2, Kdm6a and Kdm6b* RNA levels in young and old livers. Data are summarized as mean ± S.E.M. (n=8 biological replicates per group). **(B)** Western blot of H3K27me3, H3, EZH1, EZH2, KDM6A and KDM6B protein levels in young and old livers. (n=3 biological replicates per group). **(C)** Western blot of H3K27me3, H3, EZH1 and EZH2 in proliferating and quiescent cells over 14 days. Quiescence was induced by contact inhibition (n=2 biological replicates per group). **(D)** Heatmap showing EZH2 signal at differential peaks from Fig. 1G. **(E)** Genome browser snapshots of EZH2 enrichment over 4 peak regions from (D). **(F)** Western blot of H3K27me3, H3, EZH1 and EZH2 from different salt fractions of chromatin from Fig. 2B in young and old livers. On the right is the Ponceau S-stained membrane from the same experiment.

We reasoned that this switch towards EZH1 occurs due to hyper-quiescence i.e., a state of deep quiescence achieved in aged solid tissues due to prolonged stay in a post-mitotic state. To test this directly, we induced quiescence in Hep-G2 hepatoblastoma cell lines by contact inhibition (Fig. 4D). Contact inhibition-induced quiescence best mimics the state of cells in normal tissue homeostasis without nutrient deprivation. We saw an increase in H3K27me3 upon entry into quiescence with concomitant loss of EZH2 and gain of EZH1, a condition that is mimicked in aging (Fig. 4B-C). We thus propose that aged livers are in a hyper-quiescent state with a global gain in EZH1-mediated H3K27me3 and loss of EZH2.

We next wanted to understand the mechanism of local loss and gain of H3K27me3 at peak regions and age-domains respectively. Epigenetic Landscape In Silico deletion Analysis (LISA) predicted that the H3K27me3 peaks at promoters of developmental genes were mostly regulated by EZH2, and other polycomb group proteins (Table S4). Indeed, EZH2 ChIP-seq in old livers showed reduced binding at differentially regulated H3K27me3 peak regions (Fig. 4D-E, Table S1, n=4 biological replicates per group, QC metrics in Fig. S8A-G). From this data as well as the overall loss of EZH2 expression with age (Fig. 4B-C), we conclude that local loss of H3K27me3 is mediated by EZH2. Converse to the loss of signal at H3K27me3 peaks, age-domains gain signal. We wondered whether this gain of H3K27me3 signal at age-domains was due to EZH1 which increased in expression during aging (Fig. 4B-C). Given that commercial ChIP-grade antibodies detecting the mouse endogenous EZH1 protein are lacking, we blotted for EZH1/2 in the different salt fractions of chromatin in young and old livers from Fig. 2A-D. As expected, H3K27me3 was detected in the high salt fractions and old livers had stronger signal. Interestingly, EZH1 was detected in the high salt and pellet fraction in old and to a much lesser extent in young livers. EZH2 was detected mainly in the pellet fraction. We thus conclude that both EZH1 and EZH2 regulate age-domains, however, since EZH1 is expressed more in aging and detected over several high salt fractions in the old, it is likely to play a dominant role.

### Liver regeneration dilutes H3K27me3 and rejuvenates old tissue

We next wondered whether liver regeneration can epigenetically and functionally rejuvenate aged livers. Liver regeneration conventionally repopulates the parenchyma with new cells borne out of cell division, which is likely to disrupt the hyper-quiescent state. Furthermore, regeneration is hypothesized to reduce H3K27me3 by replication dilution (Jadhav et al., 2020).

The multi-lobular structure of the rodent liver enables removal of specific lobes (hepatectomy) without necrosis (Michalopoulos and Bhushan, 2021), allowing for the study of pure regenerative events unlike chemically-induced models of regeneration. We used the 70% partial hepatectomy (PH) model (Mitchell and Willenbring, 2008) to induce regeneration in young and old mice (n=3 biological replicates per timepoint, Table S1, Fig. S9A-C). Livers were archived as “before” and “after” resection samples; 48, 72, 96, 120, and 240 h post-surgery. The 240 h timepoint represents fully regenerated livers.

To assay regeneration at the transcriptome level, we analyzed our bulk RNA-seq libraries generated from ribo-depleted RNA. In pair-wise comparisons between old and young livers before resection, very few mRNAs were differentially regulated (Fig. S9D, Table S5). However, 48 h post-resection, there was a strong induction of 379 mRNAs associated with cell division in the young liver (Fig. 5A left, Table S5). Congruently, LISA identified that the top 3 upstream binding factors were E2F factors, known to regulate cell cycle entry into S-phase (Ren et al., 2002) (Table S6). The 177 mRNAs that were upregulated in the old livers 48 h after surgery instead were related to cell adhesion, inflammation, and response to glucose and insulin (Fig. 5A right) but unrelated to cell proliferation. In contrast, at the later 96 h timepoint, 199 mRNAs were upregulated in the old livers and were associated with cell division (Fig. 5B right). Sixty of the 199 mRNAs overlapped with the set of 379 mRNAs upregulated at 48 h in young livers (*p-* value=1.04e-65, representation factor 23.9; hypergeometric test). Contrary to the robust response noted in young livers at 48 h, the response in old livers at 96 h was modest with higher *p-*values and lower gene counts. Additionally, unlike the narrow time window of response in the young (48 h), the proliferative response in old livers occupied a broader window spreading from 96 to 120 h post-resection (Figs. 5B and S9E). The top candidate factor in LISA analysis for mRNAs upregulated at 96 h (Table S7) and 120 h (Table S8) in the old was also an E2F factor, E2F4. The 80 mRNAs that were upregulated in the young at 96 h were related to circadian regulation (Fig. 5B left), suggesting a return to homeostasis and regularization of normal liver functions. Of note, the 72 h and 240 h timepoints did not show many differentially expressed mRNAs (Fig. S9F-G). The 72 h timepoint likely represents a quiet phase when the young liver has mostly completed its proliferation, but the old livers have not yet induced cell cycle. In contrast, the 240 h timepoint may represent a return to quiescence for both young and old livers. We validated the RNA-seq results by tracing the cell proliferation marker Ki67 by immunohistochemistry (IHC) (Figs. S9H and S10A), IF (Figs. S9I and S10B) and RT-qPCR analyses (Fig. 5C) across the regeneration time-course. We further used RT-qPCR to measure the expression of additional cell cycle markers (Fig. 5C). All proliferation markers peaked at 48 h post-resection in young animals, while those in old animals showed an attenuated peak at ∼72-96 h. Fig. 5D summarizes the dynamics of cell proliferation mRNAs across the regeneration time-course in young and old livers. These findings were validated by an independent method, ImpulseDE2 (Fischer et al., 2018), designed to capture unimodal impulse-like patterns in high throughput time series datasets such as bulk RNA-seq. The algorithm captured the transient increase in cell proliferation genes at 48 h in the young liver followed by prompt recovery to basal levels (Fig. S9J-L, Table S9). Consistent with previous reports (Bucher and Glinos, 1950; Bucher et al., 1964; Fry et al., 1984), our results showed that old mice have a delayed and subdued regenerative response to injury.

**Fig. 5:**
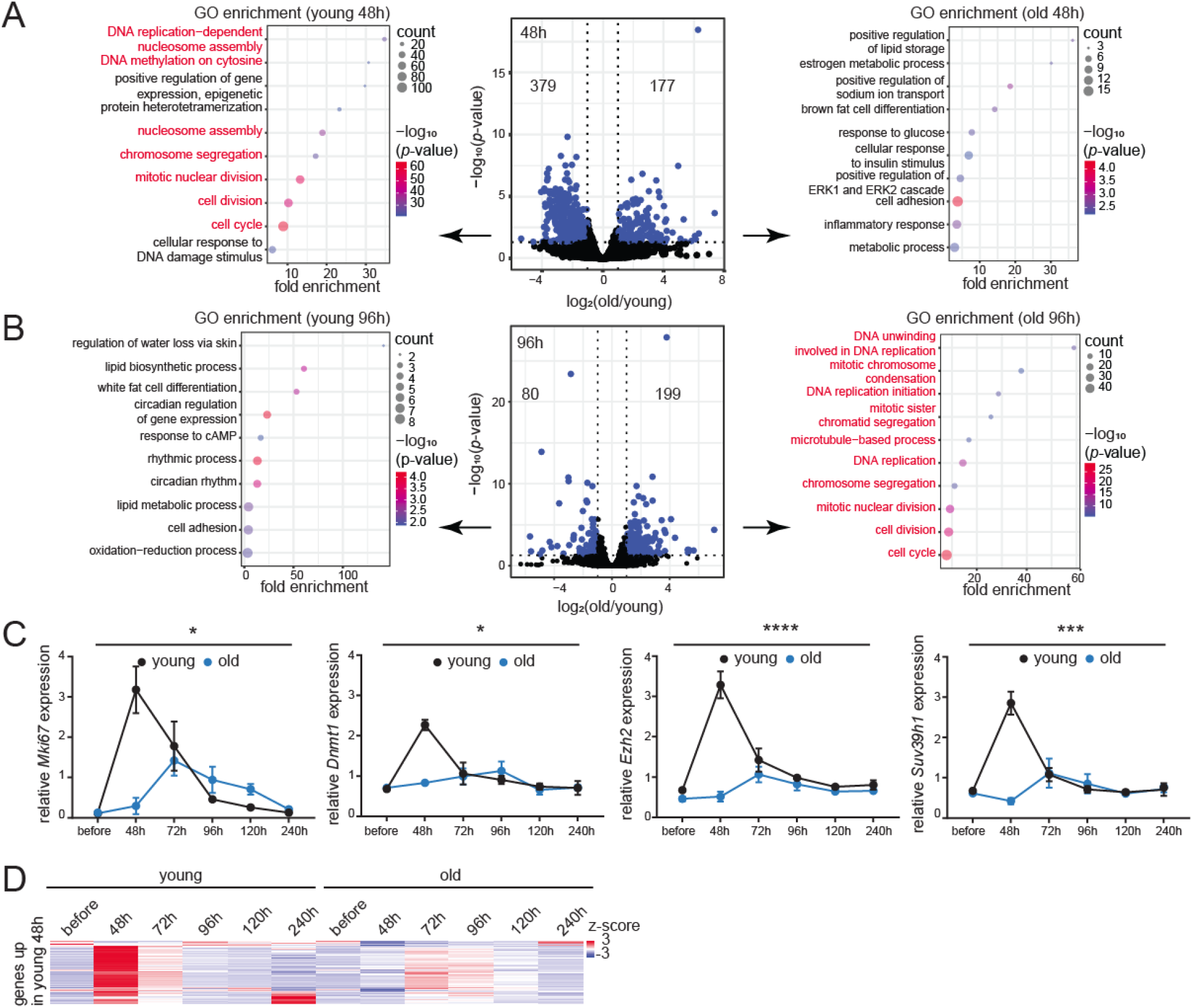
Transcriptomic signatures of aging and regeneration. **(A)** Volcano plot of differentially expressed mRNAs 48 h post-resection in young and old livers, blue dots are significant (p<0.05) mRNAs. Biological process GO terms are indicated for genes downregulated (left) or upregulated (right) in the old. Cell proliferation genes are indicated in red. **(B)** Same as (A) except samples are 96 h post-resection. **(C)** qPCR analysis of *Mki67, Dnmt1, Ezh2* and *Suv39h1* expression relative to *Actb* across the regeneration time-course. Data are summarized as mean ± S.E.M. (n=3 biological replicates per group per timepoint). * p<0.05, *** p<0.001 and **** p<0.0001 from a two-way ANOVA. **(D)** Heatmap of cell proliferation gene counts across the regeneration time-course.

Despite a reduction in regenerative response, a proportion of old hepatocytes entered cell cycle (Figs. S9H-I and S10). We reasoned that this mild proliferative event in old livers would be sufficient to dilute H3K27me3 and allow us to test if epigenomic rejuvenation reversed aspects of aging. Indeed, old regenerated livers (240 h post-surgery, Fig, 6A) showed reduction in H3K27me3 signal after regeneration (Fig. 6B, compare before and after regeneration samples). We also noted that the differences in expression of EZH1/2 between young and old livers (Fig. 4B) were abolished post-regeneration (Fig. 6C). We confirmed that regeneration reversed multiple molecular aspects of aging. For example, in regenerated livers, there was an increase in H3K27me3 at developmental gene promoters by ChIP-seq (Fig. 6D) which consequentially decreased expression of developmental genes (Fig. 6E) restoring cell identity. Importantly, we observed there was a marked reduction in age-domains (Fig. 6F). Only the global transcriptome suppression failed to be reversed (data not shown), likely due to the fact that age-domains though reduced, still persist post-regeneration (Fig. 6F). Nonetheless, PCA plots of both H3K27me3 patterns (Fig. 6G) and transcriptome (Fig. 6H) confirmed that the old regenerated livers shifted to more youthful states. To determine restoration of youthful function in old regenerated livers, we analyzed the expression profile of a group of liver-enriched genes (Table S10) obtained from the Human Protein Atlas and defined as those with higher mRNA fold compared to all other tissues (Uhlen et al., 2015). Liver-enriched genes in old regenerated livers showed expression patterns that were similar to young livers (Fig. 6I). Furthermore, we investigated if the genes that were upregulated with age (Fig. S9D, Table S5) were downregulated by regeneration. Indeed, for a major subset of genes, this was true (Fig. 6J). Taken together, these data show that liver regeneration and the ensuing replication dilution of H3K27me3, reverses multiple aspects of aging at the molecular and physiological level.

**Fig. 6:**
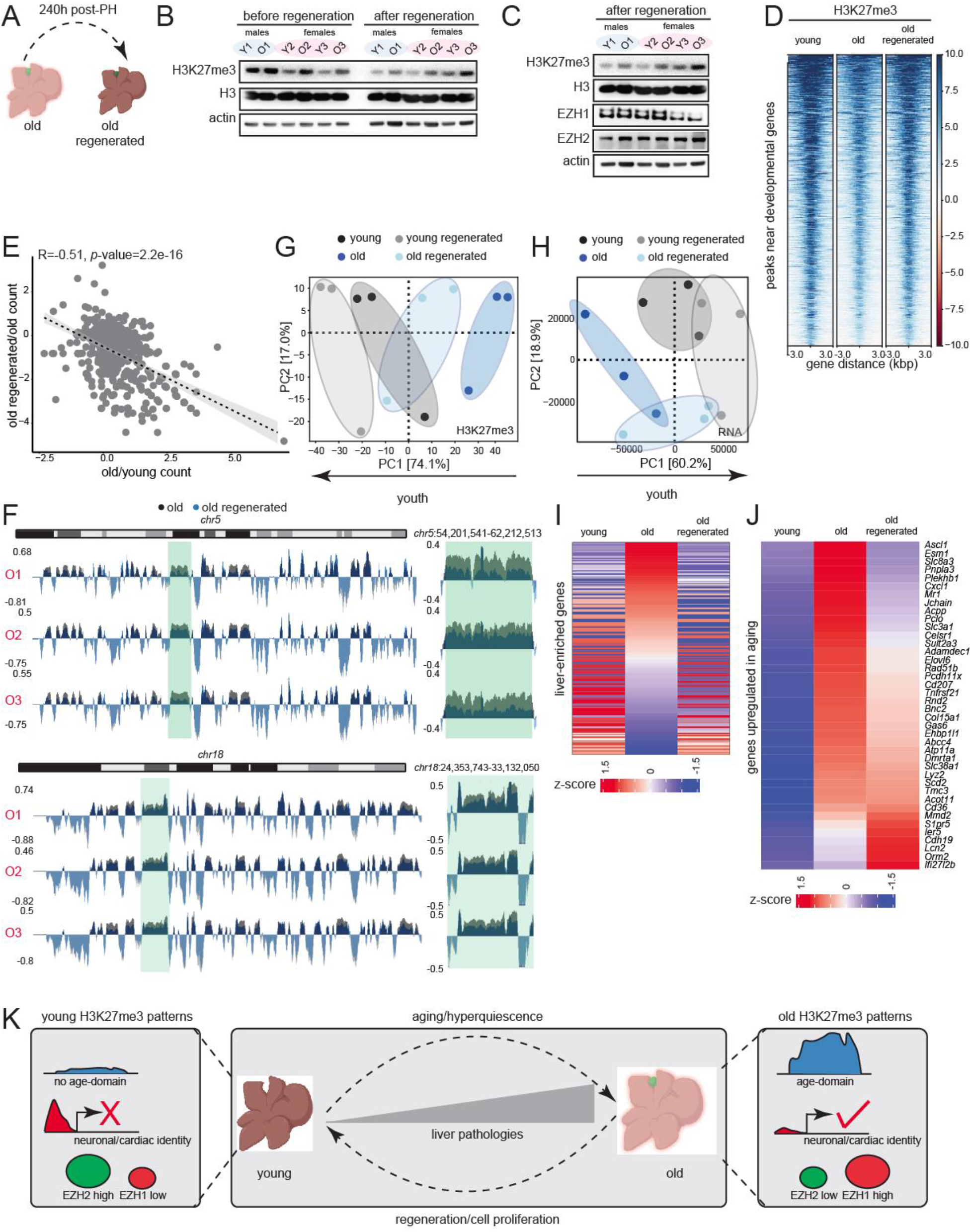
Liver regeneration dilutes H3K27me3 and rejuvenates old tissue. **(A)** Schematic of complete liver regeneration. **(B)** Western blot of H3K27me3, H3 and β-actin showing replication dilution of H3K27me3 post-regeneration. The “before regeneration” H3K27me3 blot is same as in Fig. 4B. **(C)** Western blot of H3K27me3, H3, EZH1, EZH2 and β-actin post-regeneration. For (B-C) n=3 and 2 replicates per group respectively. The “after regeneration” H3K27me3 blot is same as in (B). **(D)** Heatmap of H3K27me3 signal at differential peaks from Fig. 1G before and after regeneration. **(E)** Negative correlation of gene expression between old vs young and old regenerated vs old. The gene set corresponds to common genes that were de-repressed in aging and re-repressed post-regeneration. **(F)** Genome browser shot of overlayed H3K27me3 ChIP signal over chr5 (top) and chr18 (bottom) before and after regeneration. Green area is expanded on the right of each chromosome. **(G)** PCA plot of H3 subtracted H3K27me3 genome coverage from young, young regenerated, old, and old regenerated livers. **(H)** Same as (G) except PCA is from RNA-seq data. For (G-H) n=3 replicates per group. **(I)** Heatmap of liver-enriched gene counts in young, old, and old regenerated livers sorted on the old sample. **(J)** Heatmap of age-upregulated (from Fig. S9D) gene counts in young, old, and old regenerated livers sorted on the old sample. **(K)** Overview of H3K27me3 changes in aging and reversal by regeneration.

### Multiple tissues show features of hyper-quiescent chromatin during aging

Finally, we investigated if age-domains were evident in aged quiescent tissues other than the liver thus presenting a common epigenetic mechanism of tissue aging. We performed ChIP-seq of H3K27me3 in young and old kidney (QC metrics in Fig. S11A-G). As in the aged liver, we observed age-domains of H3K27me3 across chromosomes in the aged kidney (overlayed RPKM normalized H3K27me3 signal over chr5 and 18 shown as examples in Fig. S12A-C). Additionally, we mined published H3K27me3 ChIP-seq data from aged mouse heart and quadriceps muscle (Bou Sleiman et al., 2020) and processed the data using our ChIP-seq pipeline. The results showed evidence of H3K27me3 age-domains as in liver (Fig. S12D-E). Taken together, our results indicate that accumulation of H3K27me3 across large regions of the genome might be a universal molecular phenotype of aged quiescent tissues.

## Discussion

In this study, we carefully dissected the fundamental properties of aged chromatin using multiomics methodologies and direct imaging. In the quiescent liver, at any given time over the course of life, only 0.1% of hepatocytes are in mitosis. In general, hepatocytes stay viable and non-mitotic over a long lifespan (200-400 days) which in turn, abrogates the need for *bona fide* stem cells in the liver that are otherwise required in tissues with rapid turnover such as the gut (Magami et al., 2002). During the long post-mitotic phase of aged hepatocytes, we noted an interesting paralog switching event in the enzymes catalyzing H3K27 methylation (Fig. 4B). Notably, there was a decline in EZH2 expression with age which coincided with the age-related loss of H3K27me3 and EZH2 peaks at developmental gene promoters (Figs. 1G, 3C, 4D-E). Conversely, EZH1 protein levels increased (Fig. 4B) and co-eluted with age-domains of H3K27me3 along with remnant EZH2 (Fig. 4F). Histological and biochemical experiments have previously reported that EZH1 is expressed more in post-mitotic cells, has weaker methyltransferase activity, and represses transcription both *in vitro* and *in vivo (Margueron et al*., *2008)*. EZH1 is also reported to maintain a slow cycling undifferentiated state conferring protection against senescence in aged hematopoietic stem cells (Hidalgo et al., 2012). Additionally, H3K27me3 is known to deposit very slowly on newly replicated DNA (Alabert et al., 2015; Reveron-Gomez et al., 2018) and is maintained more on H3.3, which accumulates with age (Fig. S1A) (Kori et al., 2021). These properties of H3K27me3 and EZH1 perfectly agree with our observations of EZH1 expression, gradual H3K27me3 accumulation and global transcription suppression with age. Quiescence cultures induced by contact inhibition were able to model the accumulation of H3K27me3 and the antagonistic behavior of EZH1 and EZH2 observed in aging (Fig. 4C). We thus propose that aged cells are in a state of hyper-quiescence, i.e., they have been quiescent for a long time.

Of note, our work identifies a new H3K27me3 enrichment pattern in aged tissues, megabase-scale heterochromatin domains which we call age-domains (Figs. 1J, 2B, 2D, S3, and S4) that has not been reported before. These age-domains are not identified by traditional ChIP-seq peak calling algorithms that scan tags in short genomic regions but rather visible in chromosome-wide views. Additionally, age-domains cannot be identified in CUT&Tag type experiments that use antibody-tethered genome cutting enzymes without sonication and efficient exposure of epitopes in highly compacted chromatin. Given the scale of these enrichments, it is likely that age-domains represent the inactive B compartments in the 3D genome or lamina-associated domains (LADs). Indeed, we noted that age-domains were relatively gene poor, a feature of B compartments and LADs (van Steensel and Belmont, 2017).

The liver has remarkable regenerative capacity. In the face of injury, differentiated hepatocytes enter cell cycle and repopulate the liver with new cells although fraction of cells entering cell cycle declines with age. Knowing that H3K27me3 can be reduced by replication dilution (Jadhav et al., 2020), we inspected the old regenerated liver (240 h post-surgery) for signs of epigenetic rejuvenation. Importantly, at 240 h post-regeneration, the liver was fully quiescent representing a state where some cells have reduced their “post-mitotic age”. We found that even this limited proliferation was able to abolish differences in EZH1/2 expression (Fig. 6C), increase H3K27me3 signals at developmental genes (Fig. 6D), silence these genes (Fig. 6E), reduce age-domains (Fig. 6F) and correct age-related transcriptomic changes (Fig. 6I-J). Overall, our results clearly demonstrate that removing H3K27me3 by controlled cell division can reset the epigenetic landscape and rejuvenate old tissue. For the same reasons, age-domains are unlikely to be seen in actively dividing cells in the body such as aged blood cells or stem cells.

We further show that many organs remain in a repressed hyper-quiescent state *in vivo* and form age-domains (Figs. S11 and S12). Keeping tissues in a repressed, non-dividing state, may confer anti-tumor properties. Our study highlights the importance of monitoring chromatin in its native *in vivo* context as growth factor stimulation in cell culture may prompt cell cycle entry and replication dilution of epigenetic marks such as H3K27me3.

Based on our observations, we further speculate that some of the benefits of interventions such as Yamanaka factor-based reprogramming or parabiosis could be partially due to cell proliferation although they involve additional dedifferentiation events. Curiously, several reprogramming strategies have reported proliferation events in reprogrammed tissues (Chondronasiou et al., 2022; Ocampo et al., 2016; Sarkar et al., 2020). Similarly, heterochronic parabiosis has reported increased cell proliferation in the aged partner with reduction in repressive complexes known to inhibit proliferation (Conboy et al., 2005). Our study thus sheds light on a mechanistic basis for rejuvenation, i.e., dilution of H3K27me3.

Independently, our study makes an intriguing suggestion that either EZH2 overexpression or EZH1 inhibition may have tissue rejuvenation properties although it remains to be tested. Unfortunately, most inhibitors targeting H3K27me3 methylases are either EZH2-specific or EZH1/2 dual inhibitors. Given the high sequence similarity between EZH1 and EZH2 SET domains, design of specific inhibitors is challenging, albeit not impossible. In the absence of an EZH1-targeting drug, our study suggests cell proliferation-mediated global reduction in H3K27me3 levels could be a potential therapeutic strategy to ameliorate age-relate decline and disease.

## Supporting information

Yang_combined_supp

## Acknowledgments

We wish to acknowledge the National Institute on Aging Intramural Research Program (NIA IRP), National Institutes of Health (NIH), for financial support. We thank the Comparative Medicine Section at NIA for providing support with animal experiments. We thank Steven Henikoff for CUT&Tag-related troubleshooting. We thank Aubrey Mwinyogle and Artem Shmelev for help with the acquisition of the human-liver biopsies. We appreciate Myriam Gorospe, Weidong Wang, Ranjan Sen, Michael Seidman and Shelley Berger for critical feedback. We thank Elin Lehrmann for help with GEO upload. This work utilized the computational resources of the NIH HPC Biowulf cluster (http://hpc.nih.gov). Some figures were made using BioRender. The work was funded by NIH ZIA AG000679 to P.S., NIH ZIA BC011206 to YD, NIH S10 OD030286-01, NIH P30 CA01333047, Leukemia Research Foundation, AFAR, The Japan Agency for Medical Research and Development, Merck, and Deerfield to S.S. (Stransky) and S.S. (Sidoli).

## Author contributions

Conceptualization: NY and PS

Writing – original draft: NY and PS

Writing – review & editing: SCC and ES

Wet lab investigations: NY, LW, CYS

Bioinformatics: NY, JO, PS

Salt fractionation and AFM: DPM and YD

Histone proteomics and data analysis: SS (Stransky) and SS (Sidoli)

IHC/IF assistance: MED and JME

TEM and data analysis: MD

Sequencing: JF and SD

Animals: RD

Human liver: ES and SCC

## Figure Titles and Legends

## STAR Methods

### RESOURCE AVAILABILITY

#### Lead contact

Further information and requests for resources and reagents should be directed to and will be fulfilled by the lead contact, Payel Sen.

#### Materials availability

This study did not generate new unique reagents. Materials used in this study are listed in the Key Resources Table.

#### Data and code availability

All genome-wide datasets have been submitted to the Gene Expression Omnibus portal (GEO: GSE185708). Raw mass spec data are deposited at chorusproject.org/1769. All code used in this study is at https://github.com/PSenlab/Yang_2022. See Key Resources Table for details.

## EXPERIMENTAL MODEL AND SUBJECT DETAILS

### Cell lines and culture conditions

Hep-G2 (human male; RRID:CVCL_0027) cells were cultured in a 37°C 5% CO_2_ and 20% O_2_ humidified incubator with Dulbecco’s Modified Eagles Medium (DMEM, Gibco) supplemented with 1% penicillin/streptomycin (Thermo Fisher) and 10% Fetal Bovine Serum (Thermo Fisher). For ChIP-Rx experiments, HeLa (human female; RRID:CVCL_0030) cells cultured in the same way as HepG2 cells were used.

### Animals

This study was approved by the Animal Care and Use Committee of the NIA in Baltimore, MD under Animal Study Protocol numbers 481-LGG-2022 and 481-LGG-2025. Young and old inbred C57BL6/JN mice of both sexes were acquired from the NIA aged rodent colony (https://ros.nia.nih.gov/) and housed in rooms that were maintained at 22.2 ± 1 °C and 30-70% humidity. Routine tests are performed to ensure that mice are pathogen-free and sentinel cages are maintained and tested according to American Association for Accreditation of Laboratory Animal Care (AAALAC) criteria. The age and sex information are available in Table S1.

### Human liver samples

Core-needle biopsies of the liver were obtained from consenting patients undergoing cholecystectomy, bariatric procedures, or gastroesophageal reflux disease with mostly normal livers. Exclusion criteria included known liver disease and elevated liver enzymes. Approval was obtained by the Institutional Review Board of Ascension Saint Agnes Hospital, Baltimore (protocol number RPN 2019-016) prior to initiation of the study. Informed consent was obtained from all patients.

## METHOD DETAILS

### Induction of quiescence

To induce quiescence by contact inhibition, cells were allowed to grow until they reached 100% confluency. The cultures were maintained for 2, 7 or 14 days with media change (with serum) every two days.

### Partial hepatectomy

70% partial hepatectomy was performed following guidelines in Mitchell et al (Mitchell and Willenbring, 2008). The removed liver lobes were archived as “before resection” samples. Following specified times after surgery, the animals were sacrificed by carbon dioxide asphyxiation and cervical dislocation. The liver was dissected and either frozen in isopentane chilled with liquid nitrogen and stored in −80°C or processed for paraffin embedding and/or OCT (Tissue-Tek) encapsulation. All surgeries were performed at approximately the same time of day to prevent confounding variables such as circadian rhythms.

### Immunohistochemistry

6 µm sections of liver were cut from a paraffin block onto positively charged glass slides. Sections were deparaffinized, rehydrated and autoclaved in citrate buffer (Thermo Fisher) for antigen retrieval. Following several washes in Tris Buffered Saline with or without 0.1% Tween-20 (TBS or TBST; Pierce), the sections were blocked for 30 min with 2.5% normal goat serum (Vector Biolabs) and incubated with primary antibody overnight at 4°C in a humidified chamber. After washing with TBST, endogenous peroxidase was blocked using the Peroxidase and Alkaline Phosphatase Blocking Reagent (Agilent Dako) and the Dual-Link Envision system (Agilent Dako) was applied for 30 min at room temperature. The sections were then incubated with chromogenic substrate DAB+ (Agilent Dako) for 2 min followed by washing in deionized water, staining with Mayer’s hematoxylin (Vector Laboratories) and treatment with Scott’s tap water (Sigma). Following further washing, the sections were dehydrated and cleared with xylene before mounting with DPX mountant (Sigma Aldrich). Images were taken on a Zeiss Axiovert 200 microscope using brightfield settings. Antibodies are listed in Key Resources Table.

### Immunofluorescence

Liver sections (10-12 µm) were cut from an OCT block onto positively charged glass slides using a cryostat. The sections were permeabilized with 0.2% Triton X-100 at room temperature for 5 min. Antigen retrieval was performed by heating to 95°C for 30 min. Blocking and primary antibody incubation was performed as in IHC. The sections were then incubated with secondary antibody conjugated to a fluorescent dye for 1 h at room temperature followed by washes with TBST and staining with 5 µg/ml DAPI for 15 min at room temperature. After two washes with TBS, the sections were mounted with Epredia Lab Vision PermaFluor Aqueous Mounting Medium (Fisher Scientific). Photographs were taken using a Zeiss LSM 710 confocal microscope. Intensities were quantified using Image J (https://imagej.nih.gov/ij/). Antibodies are listed in Key Resources Table.

### RNA isolation and RT-qPCR

RNA was isolated from frozen tissue by homogenization in Trizol followed by isopropanol precipitation. The RNA was further purified using RNeasy columns (Qiagen). An on-column DNase I digestion was performed during the purification step to remove genomic DNA. The RNA amount and integrity were confirmed using the Qubit RNA HS Assay Kit and RNA IQ Assay (Thermo Fisher) respectively. Total RNA (∼500 ng) was converted to cDNA using the High Capacity RNA to cDNA kit (Thermo Fisher) for RT-qPCR analysis. 1 µl of 1:10 dilution of cDNA was analyzed by qPCR using PowerUp SYBR Green Master Mix (Thermo Fisher) following the standard curve method on a QuantStudio 7 Flex machine (Thermo Fisher). A “minus RT” control was used to confirm removal of genomic DNA. Primers are listed in Key Resources Table.

### RNA sequencing with ERCC Ex-Fold spike-in

Total RNA (∼1 µg) was used to make RNA-seq libraries following the Zymo-Seq Ribo-free Total RNA Library Kit (Zymo Research) instructions with dual indexing. Prior to ribo-depletion, total RNA from young livers were spiked with 2 µl of a 1:100 dilution of ERCC Ex-fold Mix 1 while that from old livers were spiked with 2 µl of 1:100 dilution of ERCC Ex-fold Mix 2 (Thermo Fisher). Exfold ERCC spike-ins are provided in two mixes, Mix 1 and Mix 2, that have the same pre-formulated blend of 92 transcripts but at different amounts such that a group of transcripts in Mix 1 are always at a defined fold difference from the same group in Mix 2 (Fig. 1G). There are four such groups with log_2_(Mix1/Mix2) ratios of 2 (4X), 0 (1X), -0.58 (0.67X) and -1 (0.5X). Library quality and quantity was confirmed on a BioAnalyzer (Agilent) DNA HS chip. Equimolar amounts of each library were combined, and the pooled library was further quantified using a NEBNext Library Quant Kit (New England Biolabs). The RNA-seq libraries were subjected to two rounds of 75bp paired end sequencing on a NextSeq 550 platform using a 150-cycle kit (Illumina).

### Bottom-up nanoLC-MS/MS and data analysis

#### Histone extraction and digestion

Histone proteins were extracted from nuclei pellet as previously described by Sidoli et al (Sidoli et al., 2016) to ensure good-quality identification and quantification of single histone marks. Briefly, nuclei were isolated by douncing ∼20 mg tissue in 1ml nuclei isolation buffer (15 mM Tris–HCl pH 7.5, 15 mM NaCl, 60 mM KCl, 5 mM MgCl_2_, 1 mM CaCl_2_, 250 mM sucrose, 0.2% NP-40) supplemented with 1 mM DTT, 1X Halt protease and phosphatase inhibitor cocktail (Thermo Fisher), 1 mM PMSF and 10 mM sodium butyrate. Nuclei were pelleted by centrifugation at 700 g for 5 min. Histones were acid-extracted with chilled 0.2 M sulfuric acid (5:1, sulfuric acid:pellet) and incubated with constant rotation for 4 h at 4°C, followed by precipitation with 33% trichloroacetic acid (TCA) overnight at 4°C. Then, the supernatant was removed, and the tubes were rinsed with ice-cold acetone containing 0.1% HCl, centrifuged and rinsed again using 100% ice-cold acetone. After the final centrifugation, the supernatant was discarded, and the pellet was dried using a vacuum centrifuge. The pellet was dissolved in 50 mM ammonium bicarbonate, pH 8.0, and histones were subjected to derivatization using 5 µl of propionic anhydride and 14 µl of ammonium hydroxide (Sigma Aldrich) to balance the pH at 8.0. The mixture was incubated for 15 min and the procedure was repeated. Histones were then digested with 1 µg of sequencing grade trypsin (Promega) diluted in 50 mM ammonium bicarbonate (1:20, enzyme:sample) overnight at room temperature. Derivatization reaction was repeated to derivatize peptide N-termini. The samples were dried using a vacuum centrifuge.

#### Sample desalting

Prior to mass spectrometry analysis, samples were desalted using a 96-well plate filter (Orochem) packed with 1mg of Oasis HLB C-18 resin (Waters). Briefly, the samples were resuspended in 100 µl of 0.1% TFA and loaded onto the HLB resin, which was previously equilibrated with 100 µl of the same buffer. After washing with 100 µl of 0.1% TFA, the samples were eluted with a buffer containing 70 µl of 60% acetonitrile and 0.1% TFA and then dried in a vacuum centrifuge.

#### LC-MS/MS Acquisition and Analysis

Samples were resuspended in 10 µl of 0.1% TFA and loaded onto a Dionex RSLC Ultimate 300 (Thermo Scientific), coupled online with an Orbitrap Fusion Lumos (Thermo Scientific). Chromatographic separation was performed with a two-column system, consisting of a C-18 trap cartridge (300 µm ID, 5 mm length) and a picofrit analytical column (75 µm ID, 25 cm length) packed in-house with reversed-phase Repro-Sil Pur C18-AQ 3 µm resin. Histone peptides were separated using a 30 min gradient from 1-30% buffer B (buffer A: 0.1% formic acid, buffer B: 80% acetonitrile + 0.1% formic acid) at a flow rate of 300 nl/min. The mass spectrometer was set to acquire spectra in a data-independent acquisition (DIA) mode. Briefly, the full MS scan was set to 300-1100 m/z in the orbitrap with a resolution of 120,000 (at 200 m/z) and an AGC target of 5×10e5. MS/MS was performed in the orbitrap with sequential isolation windows of 50 m/z with an AGC target of 2×10e5 and an HCD collision energy of 30.

#### Data analysis

Histone peptides raw files were imported into EpiProfile 2.0 software (Yuan et al., 2018). From the extracted ion chromatogram, the area under the curve was obtained and used to estimate the abundance of each peptide. To calculate the relative abundance of post-translational modifications (PTMs), the sum of all the different modified forms of a histone peptide was considered as 100% and the area of the particular peptide was divided by the total area for that histone peptide in all of its modified forms. The relative ratio of two isobaric forms was estimated by averaging the ratio for each fragment ion with different mass between the two species. The resulting peptide lists generated by EpiProfile were exported to Microsoft Excel and further processed for a detailed analysis. Differences between conditions were assessed using t-test statistics; significant changes were considered at the heteroscedastic t-test *p*<0.05.

### Western blot

Tissues (∼20 mg) or cells were homogenized in RIPA buffer containing 50 mM Tris pH 7.5, 0.5 mM EDTA, 150 mM NaCl, 1% NP40, 1% SDS, supplemented with 1X Halt protease and phosphatase inhibitor cocktail (Thermo Scientific), 1 mM PMSF and 10 mM sodium butyrate. The lysates were briefly sonicated with a Bioruptor (Diagenode), and cleared by centrifugation at max speed for 10 min at 4°C. In some cases, nuclear lysates were used for EZH1 western; lysates were prepared as described previously (Sen et al., 2019). The supernatants were quantified using the BCA kit (Pierce) and ∼10-30 µg total protein subjected to electrophoresis using NuPAGE 12% Bis-Tris gel in MES buffer (Thermo Fisher). The proteins were transferred to a 0.2-micron nitrocellulose membrane using a XCell II blot module (Thermo Fisher) for 1 h at 30V surrounded by ice. Proper transfer was verified by Ponceau S staining. 5% milk in TBST was used to block the membrane at room temperature for 1 h followed by primary antibody incubation at 4°C overnight. The membrane was washed and incubated with HRP-conjugated secondary antibodies (BioRad) at room temperature for 1 h. The membrane was washed again 3 times, developed, and imaged by a KwikQuant imager (Kindle Biosciences). Intensities were quantified using Image J (https://imagej.nih.gov/ij/). Antibodies are listed in Key Resources Table.

### Transmission Electron Microscopy with immunogold labeling

#### Sample processing

Animals were perfused with freshly made EM grade 100 mM Sorensen’s phosphate buffer pH 7.4 with 2% paraformaldehyde, 1% glutaraldehyde and 5 mM MgCl_2_ at 4°C. All subsequent steps were done at 4°C until 70% ethanol dehydration. Livers were carefully dissected in fixative and cut into pieces measuring no more than 2 mm^3^. Samples were rinsed thrice in buffer containing 3% sucrose for 15 min each and then osmicated in 1% osmium tetroxide (1.5% potassium ferrocyanide reduced) in 100 mM phosphate buffer, containing 5 mM MgCl_2_ for 2 hrs on ice. Tissue was put back in phosphate/sucrose buffer overnight on a cold room rocker. Samples were rinsed thrice in 100 mM maleate buffer containing 3% sucrose for 5 min each and then en-bloc stained with 2% filtered uranyl acetate in the same buffer for 1 hr. Samples were dehydrated at 4°C up to 70% ethanol after which they were brought to room temperature and further dehydrated to 100% ethanol. Liver pieces were embedded with Eponate 12 after propylene oxide transition, and finally cured in a 60°C oven for two days.

#### Immunogold labeling

Ultra-thin [60 nm (grey)] sections were picked up on glow discharged formvar coated 200 mesh nickel grids. Sections were floated on all subsequent steps. Anti-capillary tweezers were used to transfer grids and then placed on 3% sodium meta periodate (aq) twice for 15 min each. After a 15 min rinse in distilled water, grids were floated on 10 mM citrate buffer pH 6.2 for 20 min at 95°C for antigen retrieval. After cooling down, grids were placed on 0.1 M glycine in TBS for 10 min, followed by 30 min incubation in blocking buffer (1% BSA in TBST) and primary antibody (1:100) incubation overnight. Samples with no primary antibody added served as negative controls. Next day, grids were equilibrated to room temperature for 1 hr and placed in blocking solution for 10 min, followed by a 10 min rinse in TBS. Gold conjugated secondary antibodies were diluted 1:40 (6 nm goat anti-rabbit, Jackson Immunochemicals) in TBS and grids were incubated for 2 hrs at room temperature in a humidified chamber. After a 10 min TBS incubation followed by a quick distilled water rinse, grids were hard fixed in 2% glutaraldehyde in 100 mM sodium cacodylate buffer for 5 min. After a brief distilled water rinse, grids were stained with 2% uranyl acetate in 50% methanol for 10 min, rinsed again with distilled water, blot dried and allowed to sit in grid boxes overnight before viewing. Sections were viewed on a Hitachi H 7600 TEM operating at 90 kV and digital images captured with an ER-80 (8 megapixel) CCD camera, by AMT. Antibodies are listed in Key Resources Table.

#### Immunogold particle quantification

Ten cells were randomly chosen from each sample (60 nm sections). In each cell, 5 non-overlapping regions of interest (ROI) in the nucleus were identified. For each ROI, the following were counted: total gold number, gold number in cytoplasm, gold number at nuclear periphery (within 200 nm), and gold number in nuclear matrix. Clusters (≥3 gold particles) in the nuclear matrix, nuclear periphery and cytoplasm for each ROI were also quantified.

### Chromatin immunoprecipitation and sequencing

Crude nuclei preparations were made from ∼100 mg frozen tissue by douncing in nuclei preparation buffer (10 mM Tris pH 7.4, 10 mM NaCl, 3 mM MgCl_2_, 0.1% Tween 20, 0.1% NP-40, 0.01% digitonin and 1 mM BSA, supplemented with protease inhibitors and 1 mM sodium butyrate). After centrifugation, the nuclei pellet was crosslinked with 1% formaldehyde in 2 ml PBS by constant rotation at room temperature for 10 min followed by quenching with 125 mM glycine and two PBS washes. The nuclei were lysed with nuclei lysis buffer (10 mM Tris-HCl pH 7.4, 100 mM NaCl, 1 mM EDTA, 0.5 mM EGTA, 0.1% sodium-deoxycholate, 0.5% N-lauroylsarcosine supplemented with protease inhibitors and 1mM sodium butyrate) and sheared to <500bp using a Covaris S220 Ultrasonicator. An aliquot of the sheared chromatin sample was removed to check sonication success and to quantify the amount of chromatin. The immunoprecipitation, wash and elution steps were performed as reported previously (Sen et al., 2019) except ∼4% HeLa chromatin was spiked in to 2 µg mouse chromatin prior to immunoprecipitation (only for liver samples). EZH2 ChIP was performed using the SimpleChIP plus sonication kit (CST #56383). ChIP enrichment was verified by qPCR around the p16 promoter (−200bp, -1000bp and -5000bp upstream of TSS for H3K27me3 or around the *HoxA1* and *HoxD10* locus for EZH2), actin promoter as the negative control locus and IgG as specificity control for the antibody. DNA (∼5 ng) was used to prepare libraries with the NEBNext Ultra II library preparation kit with unique dual index primers (New England Biolabs). The library quality and quantity were verified by BioAnalyzer DNA 1000 (Agilent) run and qPCR with NEBNext Library Quant kit (New England Biolabs) respectively. The liver libraries were pooled and paired-end sequenced on the NextSeq 2000 platform (Illumina) using the P2 or P3 100 cycle kit. Antibodies are listed in Key Resources Table.

### Cut&Tag

Cut&Tag was performed following guidelines in the CUT&Tag@home v.1 protocol by Henikoff et al (Henikoff et al., 2020). Briefly, nuclei were isolated from ∼20mg frozen human liver tissue using Minute Detergent-free Nuclei Isolation Kit (Invent Biotechnologies). The nuclei were permeabilized with 0.1% Triton X-100 and their integrity and number was checked under a microscope using a hemocytometer. ∼10,000 nuclei per sample were bound to Concanavalin A beads (Bangs Laboratories). The bead-bound nuclei were incubated with primary antibody (1 h at room temperature), secondary antibody (0.5 h at room temperature) and home-made pA-Tn5 transposome (1 h at room temperature). Targeted tagmentation was then performed by addition of buffer containing MgCl_2_ for 1 h at 37°C. The tagmented DNA was washed with a TAPS-EDTA wash buffer (10 mM TAPS, pH 8.5, 0.2 mM EDTA) to remove excess salt and Mg^2+^ ions followed by targeted release with 0.1% SDS and neutralization of SDS by Triton X-100. The released DNA was amplified using dual-indexing primers following a PCR protocol that biases amplification of short DNA fragments. The excess primers were removed by a 1.3X SPRI bead purification (Beckman Coulter). Library quality and quantity was confirmed on a BioAnalyzer (Agilent) DNA HS chip. Equimolar amounts of each library were combined, and the pooled library was further quantified using a NEBNext Library Quant Kit (New England Biolabs). The CUT&Tag libraries were subjected to 50bp paired end sequencing on a NextSeq 2000 platform using a P2 100-cycle kit (Illumina). Antibodies are listed in Key Resources Table.

### Salt fractionation of chromatin

Nuclei was released from frozen tissue by douncing in TM2 buffer (10 mM Tris HCl, pH 7.4, 2 mM MgCl_2_ supplemented with 1X Halt protease and phosphatase inhibitor and 0.5 mM PMSF), pelleted by centrifugation and washed once to remove debris. The washed pellet was resuspended in TM2 buffer with 1 mM CaCl_2_ and 12000 U of MNase (New England Biolabs), incubated at 23°C for 15 minutes and the reactions stopped by addition of 0.5 mM EGTA. Approximately 30% was saved for analysis (“0”) and the rest was pelleted by centrifugation. The supernatant was removed and saved as “Supernatant” fraction after clearing. The nuclei were washed once with TM2 buffer and then resuspended in 70 µl Triton buffer (10 mM Tris–HCl pH 7.4, 2 mM MgCl_2_, 2 mM EGTA, 0.1% Triton X-100 supplemented with 1X Halt protease and phosphatase inhibitor and 0.5 mM PMSF) and evenly divided. The first aliquot was saved as “Input” while the second was used for sequential salt extraction by using Triton buffer with either 67.5, 150, 250 or 350 mM NaCl. Each extraction was done by incubating the resuspended pellet at 4°C for 2 hours. After each extraction, the nuclei were pelleted by centrifugation and the supernatant saved as “67.5”, “150”, “250” or “350” mM fraction. Finally, the remainder pellet which corresponds to ∼5–10% of chromatin, was resuspended in 35 µl of TNE buffer (10 mM Tris–HCl pH 7.4, 200 mM NaCl, 1 mM EDTA supplemented with 1X Halt protease and phosphatase inhibitor) and labeled as “pellet” fraction. All extractions were performed under physiological concentrations of Mg^2+^ to preserve nuclear and chromatin integrity. DNA was purified from each of these fractions using DNA clean and concentrator columns (Zymo Research). Purified DNA (∼10 ng) was used to make libraries using the NEBNext Ultra II kit (New England Biolabs) and sequenced on the NextSeq 2000 (Illumina) with a 51PE format.

### Atomic Force Microscopy

Nuclei was extracted from livers (Nabbi and Riabowol, 2015) and chromatin was digested with 1 U MNase (Sigma) for 2 minutes in 2 ml 0.1 M TE (10 mM Tris pH 6.8, 0.2 mM EDTA, 100 mM NaCl) supplemented with 1.5 mM CaCl_2_. MNase reactions were quenched with 10 mM EGTA and centrifuged at 94 g at 4°C. Supernatant was removed and chromatin was extracted overnight at 4°C in 0.5X PBS (67.5 mM NaCl) with a protease inhibitor cocktail (Roche) on an end-over-end rotator. Extracted chromatin was subsequently diluted 800-fold to create a single layer of chromatin when deposited on mica. Mica was freshly cleaved and functionalized with aminopropyl-silantrane (APS) before chromatin was deposited (Walkiewicz et al., 2014). Images were obtained with commercial AFM equipment (Oxford Instruments, Asylum Research’s Cypher S AFM) with silicon cantilevers (OTESPA-R3 from Olympus with nominal resonances of ∼300 kHz, stiffness of ∼42 N/m) in non-contact tapping mode in air. Images were processed as previously reported (Walkiewicz et al., 2014). Image analysis was performed using Image J (https://imagej.nih.gov/ij/) and statistical analysis and visualization were done in R/4.1.1.

### MNase titration

An MNase titration was performed with 20 mg frozen liver tissue. Briefly, the frozen tissue was ground to a fine powder using a liquid nitrogen cooled mini mortar and pestle set (Bel-Art). The ground tissue was crosslinked first with 3 mM DSG and then with 1% formaldehyde, each for 10 min at room temperature. The crosslinked sample was centrifuged, and the pellet washed twice with wash buffer and sequentially filtered through 200 µm and 50 µm filters. Following centrifugation, the pellet was resuspended in nuclease digestion buffer with MNase and incubated at 22°C for exactly 15 min. The reaction was stopped by addition of 50 mM EGTA and 1% SDS. To check MNase digestion, ∼2.5 µl of the sample was treated with proteinase K, reverse crosslinked and purified using the DNA Clean and Concentrator kit (Zymo Research). The fragment size distribution was checked on a BioAnalyzer using a DNA HS kit (Agilent).

### Bioinformatic analysis

#### RNA-seq analysis

Illumina sequencing reads (∼32 million paired-end reads per sample) were de-multiplexed using bcl2fastq/2.20.0. Reads were trimmed to remove adapter sequences using trimmomatic/0.39 (Bolger et al., 2014). The quality of the resulting FASTQs was assessed using FASTQC/0.11.9(Andrews, 2017) and reads were aligned to the mouse reference genome (assembly GRCm38/mm10, concatenated with the ERCC transcripts) using STAR/2.7.5b (Dobin et al., 2013). BAM files were sorted and indexed using samtools/1.10 (Li et al., 2009), and duplicates were removed using picard/2.20.8. The BAM files were then filtered to retain alignments with a minimum mapping quality of 10 using samtools/1.10 (Li et al., 2009). The featureCounts function of the Rsubread R package/2.6.4 (Liao et al., 2019) was used to estimate counts for all mRNA and ERCC transcripts.

Counts for the ERCC transcripts were RPKM normalized then filtered for 1 RPKM. ERCC transcripts were then plotted against their known molecular concentration. Sex- and time-matched fold-change ratios (young vs old) were calculated for each ERCC transcript and plotted against their known fold-change ratio. Linear regression analyses were performed to assess dose-response relationships using GraphPad Prism/9.0.0 (121).

Differential gene expression analysis (excluding ERCC transcripts) between old and young samples was performed separately for each timepoint using the R Bioconductor package, DESeq2/1.30.1 (Love et al., 2014). Additionally, since pairwise comparison approaches may fail to account for temporal dependencies in time-course experiments(Yuan and Kendziorski, 2006), the ImpulseDE2 R package/0.99.10 (Fischer et al., 2018) was used to also assess transient (temporal) and monotonous (permanent) transcriptional changes occurring during the liver regenerative process. Heatmaps of the impulse-fitted data by time point were created using the R Bioconductor package, ComplexHeatmap/2.6.2 (Gu et al., 2016). Both analyses were conducted at a Benjamini-Hochberg corrected *p*-value (false discovery rate, FDR) significance threshold of 0.05 (Benjamini and Hochberg, 1995).

Comparisons between RNA-seq and ChIP-seq data was performed by correlating the fold-change ratio in gene expression (old vs young) with the fold-change ratio in enrichment (old vs young) for the corresponding gene.

#### ChIP-seq analysis

Illumina sequencing reads (∼50 million paired end reads per sample) were de-multiplexed generating compressed FASTQ files by the on-board DRAGEN informatics pipeline (Illumina DRAGEN FASTQ Generation – 3.7.4) on the NextSeq 2000. The FASTQ files were trimmed to remove adapter sequences with trimgalore/0.6.6 and the qualities of the FASTQs assessed using FASTQC/0.11.9 (Andrews, 2017). The reads were aligned to the GRCm38/mm10 and GRCh38/hg38 genome assemblies using bowtie/2-2.4.2 end-to-end parameter. Sam output files were then filtered to retain alignments with a minimum mapping quality of 10 using samtools/1.9 (Li et al., 2009) and PCR duplicates were removed using picard/2.23.7. Sambamba/0.7.1 (Tarasov et al., 2015) was used to retain only uniquely mapping reads. Reads mapping to the Encyclopedia of DNA Elements (ENCODE) blacklisted regions were also removed from the analysis. There were no statistically significant differences in sequencing depth, alignment rate or alignable fragments per million across the young and old groups in any tissue. The bamCoverage function in deepTools/3.5.0 (Ramirez et al., 2014) was used to generate RPKM normalized bigWig files. H3 and input reads were subtracted from H3K27me3 and IgG respectively with bigwigCompare. Peaks were called using peakranger/1.18 with the bcp parameter using H3 and input as background for H3K27me3 and IgG respectively from the same animal. Differential peak analysis was performed using DiffBind/3.2.6 (Stark and Brown, 2011).

Scale factor for spike-in normalization was calculated as follows. For each sample, *a* is defined as the scale factor, *b* is the percentage mapped to GRCm38/mm10 genome in input, and *c* is the percentage mapped to GRCh38/hg38 genome (spike-in). Based on our experimental design, the fraction of the spike-in human genome over the total mouse genome in each sample should be the same as in input and similar across all samples.

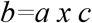

or,

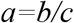

For visualization, bigWig files were generated using deepTools/3.5.0 bamCoverage with the scale parameter set as the calculated scale factor *a*1000* for each sample.

#### CUT&Tag analysis

CUT&Tag analysis with *E. coli* spike-in normalization was performed as outlined in Zheng et al, following the protocols.io tutorial (https://dx.doi.org/10.17504/protocols.io.bjk2kkye). Briefly, Illumina sequencing reads (∼6 million paired end reads per sample) were de-multiplexed generating compressed FASTQ files by the on-board DRAGEN informatics pipeline (Illumina DRAGEN FASTQ Generation – 3.7.4) on the NextSeq 2000. The qualities of the FASTQs were assessed using FASTQC/0.11.9 (Andrews, 2017). The reads were aligned to the GRCh38/hg38 genome with bowtie/2-2.4.2 using the end-to-end parameter. Sam output files were then filtered to retain alignments with a minimum mapping quality of 2 using samtools/1.9 (Li et al., 2009). Reads mapping to Encyclopedia of DNA Elements (ENCODE) blacklist regions were removed from the analysis. There were no statistically significant differences in sequencing depth, alignment rate or alignable fragments per million across the young and old groups in any tissue. The bamCoverage function in deepTools/3.5.0 was used to generate RPKM (reads per kilobase per million mapped reads) normalized bigWig files. H3 was subtracted from H3K27me3 and IgG with bigwigCompare function of deepTools/3.5.0. Differentially enriched sites were identified using the R Bioconductor package diffReps/1.55.6 (Shen et al., 2013).

#### Salt fractionation analysis

Illumina sequencing reads (∼54 million paired end reads per sample) were de-multiplexed generating compressed FASTQ files by the on-board DRAGEN informatics pipeline (Illumina DRAGEN FASTQ Generation – 3.7.4) on the NextSeq 2000. The FASTQ files were trimmed to remove adapter sequences with cutadapt/3.0 and the qualities of the FASTQs assessed using FASTQC/0.11.9 (Andrews, 2017). The reads were aligned to the GRCm38/mm10 genome using bowtie/2-2.4.2 using the end-to-end parameter. Sam output files were then filtered to keep alignments with a minimum mapping quality of 10 using samtools/1.13 (Li et al., 2009) and PCR duplicates removed with picard/2.23.7. Sambamba/0.7.1 was used to retain only uniquely aligned reads. Reads mapping to the Encyclopedia of DNA Elements (ENCODE) blacklisted regions were also removed from the analysis. There were no statistically significant differences in sequencing depth, alignment rate or alignable fragments per million across the young and old groups in any tissue. The bamCoverage function in deepTools/3.5.1 was used to generate RPKM (reads per kilobase per million mapped reads) normalized bigWig files. Input reads were subtracted from the different salt fractions and pellet with bigwigCompare.

#### PCA plots

RNA-seq PCA plot was generated in R/4.0 with the DESeq2 output and plotted with ggplot2. ChIP-seq peak PCA was generated with DiffBind (Stark and Brown, 2011).

#### Area under the curve (AUC) calculation

bwtool/1.0 (Pohl and Beato, 2014) was used to get genome coverage information (AUC) across regions of interest and then plotted with ggplot2 in R/4.0.

#### Annotation

Genomics regions were annotated using ChIPseeker/1.26 annotatePeak function in R/4.0 (Yu et al., 2015).

#### Correlation matrix

Correlation matrices were generated using the plotCorrelation function of deepTools/3.5.1 using the multibigWig summary files and a Spearman correlation.

#### Heatmaps

Heatmaps were made with deepTools/3.5.0. The computeMatrix function was first used to calculate the signal intensity on the H3K27me3 differential peak regions and then heatmap drawn with the plotHeatmap function.

#### Gene Ontology (GO) analysis

GO analysis of the differentially expressed mRNAs, differential peaks or different salt fractions (old vs young) was performed using DAVID/6.8 (Huang da et al., 2009) with *Mus musculus* genes as background. The top 10-20 significant GO terms (reported as *q*-values using a Fisher’s exact test with Benjamini correction) for the Biological Process category (sorted by fold enrichment) are reported.

#### Genome browser tracks

Genome browser tracks were created for individual and pooled (across replicates) samples by converting the BAM files to bigWig files using the bamCoverage function of deepTools/3.5.0 then uploaded on the University of California, Santa Cruz Genome Browser using either custom tracks or track hubs.

#### Hypergeometric test

Hypergeometric tests were performed with the online web calculator http://nemates.org/MA/progs/overlap_stats.cgi.

## QUANTIFICATION AND STATISTICAL ANALYSIS

Statistical analyses for all experiments were performed in GraphPad Prism/9.0.0 or R Stats/4.0.5. Statistical data are presented as mean ± S.E.M or S.D. Sample size (n), statistical tests used, and p-values are specified in the figure legends.

