## Supplementary material for "A hyper-quiescent chromatin state formed during aging is reversed by regeneration": Yang_combined_supp

### Supplemental Information

### Supplementary Figure Titles and Legends

#### Fig. S1: Increased H3K27me3 is a signature of aging

**(A)** H3.3/H3 % from LC-MS/MS raw area values in young and old livers. (n=3 biological replicates per group). **(B)** Overall abundance of monomethyl (me1), dimethyl (me2), trimethyl (me3) and acetyl (ac) -ated histones from LC-MS/MS in young and old livers. Data are summarized as mean ± S.E.M. (n=3 biological replicates per group). **(C)** Volcano plot of single hPTMs from LC-MS/MS data in muscle stem cells (Schworer et al., 2016). **(D)** Same as (C) except data is from human post-mortem brain samples (Nativio et al., 2020). The significantly increased (red) and decreased (blue) modifications are labeled (two-tailed unpaired t-test). **(E)** Representative H3K27me3 IF images. Scale bar is 10 µm. H3K27me3 intensity is plotted on the right (n=3 biological replicates, 10 fields per group). * $p<0.05$ in a Welch's two-tailed unpaired t-test. **(F)** Human spike-in alignment rate plotted as a % in H3, input and H3K27me3 immunoprecipitated samples (n=3 biological replicates per group). **(G)** PCA plot of H3 subtracted H3K27me3 genome coverage from liver ChIP-seq data. **(H)** Same as (G) except PCA is from called peaks. Note for both (G) and (H), PC1 accounts for the variation in age and PC2 accounts for the variation in sex. **(I)** Bar plot of liver H3K27me3 ChIP-seq peak numbers. Data are summarized as mean ± S.E.M. (n=3 biological replicates per group). **(J)** Violin plots of H3K27me3 peak characteristics in young and old livers. **** indicates $p<0.0001$ from a Welch's two-tailed unpaired t-test.

#### Fig. S2: Quality control of mouse liver H3K27me3 ChIP-seq libraries

**(A)** Sequencing depth of H3K27me3, H3 and input ChIP libraries made from young and old mouse livers (n=3 biological replicates per group). **(B)** Number of alignable (to GRCm38/mm10) fragments obtained after sequencing H3K27me3, H3 and input ChIP libraries. **(C)** The alignment rates of the H3K27me3, H3 and IgG ChIP libraries are plotted as a percentage of mapped fragments. **(D)** The duplication rates of the sequenced libraries are shown. Duplicates were removed prior to downstream analysis. **(E)** The estimated library size (i.e., the estimated number of unique molecules) in each library is plotted. **(F)** The unique fragment number is calculated by the mapped (to GRCm38/mm10) fragment number * (1-DuplicationRate/100). **(G)** A Spearman correlation plot of H3 subtracted H3K27me3 signal across samples.

#### Fig. S3: H3K27me3 age-domains are heterochromatinized during aging (chr5)

**(A)** Schematic showing the samples and comparisons made in (B-C). **(B)** Genome browser shot of overlayed H3K27me3 ChIP signal (H3 subtracted) over chr5 in sex-matched pairs of young and old mouse livers. **(C)** Genome browser shot of H3K27me3 ChIP signal (H3 subtracted) over chr5 in individual young and old mouse livers. For (B-C), n=3 biological replicates per group and green area in the whole chromosome is expanded on the right.

#### Fig. S4: H3K27me3 age-domains are heterochromatinized during aging (chr18)

**(A)** Schematic showing the samples and comparisons made in (B-C). **(B)** Genome browser shot of overlayed H3K27me3 ChIP signal (H3 subtracted) over chr18 in sex-matched pairs of young and old mouse livers. **(C)** Genome browser shot of H3K27me3 ChIP signal (H3 subtracted) over chr18 in individual young and old mouse livers. For (B-C), n=3 biological replicates per group and green area in the whole chromosome is expanded on the right.

#### Fig. S5: Quality control and inter-fraction correlation of DNA libraries from young and old salt fractionated liver chromatin

**(A)** Sequencing depth of libraries made from salt fractionated chromatin of young and old livers (n=2 biological replicates per group). **(B)** Number of alignable (to GRCm38/mm10) fragments obtained after sequencing of the DNA libraries. **(C)** The alignment rates of the libraries from different salt fractions are plotted as a percentage of mapped fragments. **(D)** The duplication rates of the sequenced libraries are shown. Duplicates were removed prior to downstream analysis. **(E)** The estimated library size (i.e., the estimated number of unique molecules) in each library is plotted. **(F)** The unique fragment number is calculated by the mapped (to GRCm38/mm10) fragment number * (1-DuplicationRate/100). **(G)** A Spearman correlation plot of input subtracted genome coverage from the different salt fractions. Replicates were pooled.

**Fig. S6: Age-domains are heterochromatinized while H3K27me3 peak regions are euchromatinized during aging**

**(A)** Genome browser snapshot of salt fraction enrichments over chr18 in both young and old samples. Inset 1 is a region in chr18 that is euchromatinized with age (green shade) and inset 2 is a region that is heterochromatinized with age (orange shade). Note the overlap of 350 mM and pellet fractions in the old with a H3K27me3 domain in inset 2. Green and orange areas in the whole chromosome are expanded on the right.

**Fig. S7: Quality control of human liver H3K27me3 CUT&Tag libraries**

**(A)** Sequencing depth of H3K27me3 and H3 and IgG CUT&Tag libraries made from young and old human liver biopsies (n=3 young and 5 old biological replicates). **(B)** Number of alignable (to GRCh38/hg38) fragments obtained after sequencing H3K27me3, H3 and IgG CUT&Tag libraries. **(C)** The alignment rates of the H3K27me3, H3 and IgG CUT&Tag libraries are plotted as a percentage of mapped fragments. **(D)** The duplication rates of the sequenced libraries are shown. Duplicates were not removed prior to downstream analysis. **(E)** The estimated library size (i.e., the estimated number of unique molecules) in each library is plotted. **(F)** The unique fragment number is calculated by the mapped (to GRCh38/hg38) fragment number * (1-DuplicationRate/100). **(G)** A Spearman correlation plot of H3 subtracted H3K27me3 signal across samples. **(H)** Volcano plot of diffReps output showing differential H3K27me3 peaks in young and old human livers. Significant peaks are indicated in blue and correspond to those with an FDR<0.05. **(I)** Top 20 GO terms associated with genes near the 1117 young peaks identified as differentially enriched in young (H).

**Fig. S8: Quality control of mouse liver EZH2 ChIP-seq libraries**

**(A)** Sequencing depth of EZH2 and input ChIP libraries made from young and old mouse livers (n=3 biological replicates per group). **(B)** Number of alignable (to GRCm38/mm10) fragments obtained after sequencing EZH2 and input ChIP libraries. **(C)** The alignment rates of the EZH2 ChIP libraries are plotted as a percentage of mapped fragments. **(D)** The duplication rates of the sequenced libraries are shown. Duplicates were removed prior to downstream analysis. **(E)** The estimated library size (i.e., the estimated number of unique molecules) in each library is plotted. **(F)** The unique fragment number is calculated by the mapped (to GRCm38/mm10) fragment number * (1-DuplicationRate/100). **(G)** A Spearman correlation plot of input subtracted EZH2 signal across samples.

**Fig. S9: Transcriptomic signatures of aging and regeneration**

**(A)** Schematic of experimental procedure, timeline, and assays. **(B)** Steps of 70% partial hepatectomy. **(C)** Age and sex distribution of mice used. Data are summarized as mean ± S.E.M. with individual data points representing each mouse (n=10 males and 20 females). **(D)** Volcano plot of differentially expressed mRNAs in young and old livers, blue dots are significant (p<0.05) mRNAs. Biological process GO terms are indicated for genes upregulated in the old (right). **(E)** Same as (D) except in young and old livers 120 h post-resection. Cell proliferation genes in the GO analysis are in red. **(F)** Same as (D) except in 72 h post-resection. **(G)** Same as (D) except in young and old livers 240 h post-resection. For (F-G) genes were too few for GO analysis. **(H)** Quantification of % positive cells (left) and intensity (right) of Ki67 in an immunohistochemistry assay across the regeneration time-course in young and old livers. **(I)** Quantification of % positive cells (left) and intensity (right) of Ki67 in an immunofluorescence assay across the regeneration time-course in young and old livers. For (H-I), ** p<0.01, *** p<0.001 and **** p<0.0001 from a Kolmogorov-Smirnov test with corrections for multiple comparisons. An individual data point indicates one field. **(J)** Heatmap of impulse-fitted RNA-seq data by time point (ImpulseDE2 output) showing upregulated mRNAs (red) and downregulated mRNAs (blue) in young. **(K)** GO terms associated with transiently upregulated genes in the young identified in (J). Cell proliferation genes are indicated in red. **(L)** GO terms associated with transiently downregulated genes in the young identified in (J).

**Fig. S10: Delayed and reduced liver regeneration in aged mice**
**(A)** Representative images of Ki67 immunohistochemistry in young and old liver sections across the regeneration time-course. **(B)** Representative images of Ki67 immunofluorescence in young and old liver sections across the regeneration time-course. For both (A-B), scale bar is 50 µm.

**Fig. S11: Quality control of mouse kidney H3K27me3 ChIP-seq libraries**

**(A)** Sequencing depth of H3K27me3, H3 and input ChIP libraries made from young and old mouse kidneys (n=4 biological replicates per group). **(B)** Number of alignable (to GRCm38/mm10) fragments obtained after sequencing H3K27me3, H3 and IgG ChIP libraries. **(C)** The alignment rates of the H3K27me3, H3 and IgG ChIP libraries are plotted as a percentage of mapped fragments. **(D)** The duplication rates of the sequenced libraries are shown. Duplicates were removed prior to downstream analysis. **(E)** The estimated library size (i.e., the estimated number of unique molecules) in each library is plotted. **(F)** The unique fragment number is calculated by the mapped (to GRCm38/mm10) fragment number * (1-DuplicationRate/100). **(G)** A Spearman correlation plot of H3 subtracted H3K27me3 signal across samples.

**Fig. S12: Multiple tissues show features of hyper-quiescent chromatin during aging**

**(A)** Schematic showing the samples and comparisons made in (B-C). **(B)** Genome browser shot of overlayed H3K27me3 ChIP signal (H3 subtracted) over chr5 in sex-matched pairs of young and old mouse kidneys. **(C)** Genome browser shot of H3K27me3 ChIP signal (H3 subtracted) over chr18 in individual young and old mouse kidneys (n=4 biological replicates per group). Green area in the whole chromosome is expanded on the right. **(D)** Schematic of samples and genome browser shot of H3K27me3 ChIP signal (input subtracted) over chr1 and chr3 in young (6 months) and old (24 months) mouse hearts. **(E)** Schematic of samples and genome browser shot of H3K27me3 ChIP signal (input subtracted) over chr5 and chr18 in young (6 months) and old (24 months) mouse quadriceps muscle. For both (D-E), dataset was mined from Sleiman et al, 2020 (Bou Sleiman et al., 2020) and replicates were merged. Green areas show H3K27me3 age-domains.

**Supplementary Tables Titles**

**Table S1: Sample information.**

Sheet 1: RNA-seq and mass spec
Sheet 2: Transmission Electron Microscopy (TEM)
Sheet 3: ChIP-seq
Sheet 4: Salt fraction
Sheet 5: Atomic Force Microscopy (AFM)
Sheet 6: CUT&Tag
Sheet 7: Western blot
Sheet 8: qPCR

**Table S2: Mass spectrometric quantification of hPTM data from young and old livers.**

Sheet 1: full data
Sheet 2: single PTMs

**Table S3: DiffBind output of unique H3K27me3 peaks in from uninjured mouse livers.**

**Table S4: LISA output of key transcription factors and chromatin regulators acting at promoters of top**

**500 genes differentially bound by H3K27me3 in young mouse livers.**

**Table S5: List of differentially expressed genes over the regeneration time-course in young and old livers.**

Sheet 1: before surgery (old vs young)
Sheet 2: 48 h post-surgery (old vs young)
Sheet 3: 72 h post-surgery (old vs young)
Sheet 4: 96 h post-surgery (old vs young)
Sheet 5: 120 h post-surgery (old vs young)
Sheet 6: 240 h post-surgery (old vs young)

**Table S6: LISA output of key transcription factors and chromatin regulators acting at promoters of genes differentially upregulated in young livers 48 h post-resection.**

**Table S7: LISA output of key transcription factors and chromatin regulators acting at promoters of genes differentially upregulated in old livers 96 h post-resection.**

**Table S8: LISA output of key transcription factors and chromatin regulators acting at promoters of genes differentially upregulated in old livers 120 h post-resection.**

**Table S9: List of genes up- or downregulated (transiently or monotonously) during the regeneration time-course in young and old livers as measured by ImpulseDE2.**

Sheet 1: transiently up in young
Sheet 2: transiently down in young
Sheet 3: monotonously up in young
Sheet 4: monotonously down in young

**Table S10: Liver-enriched genes**

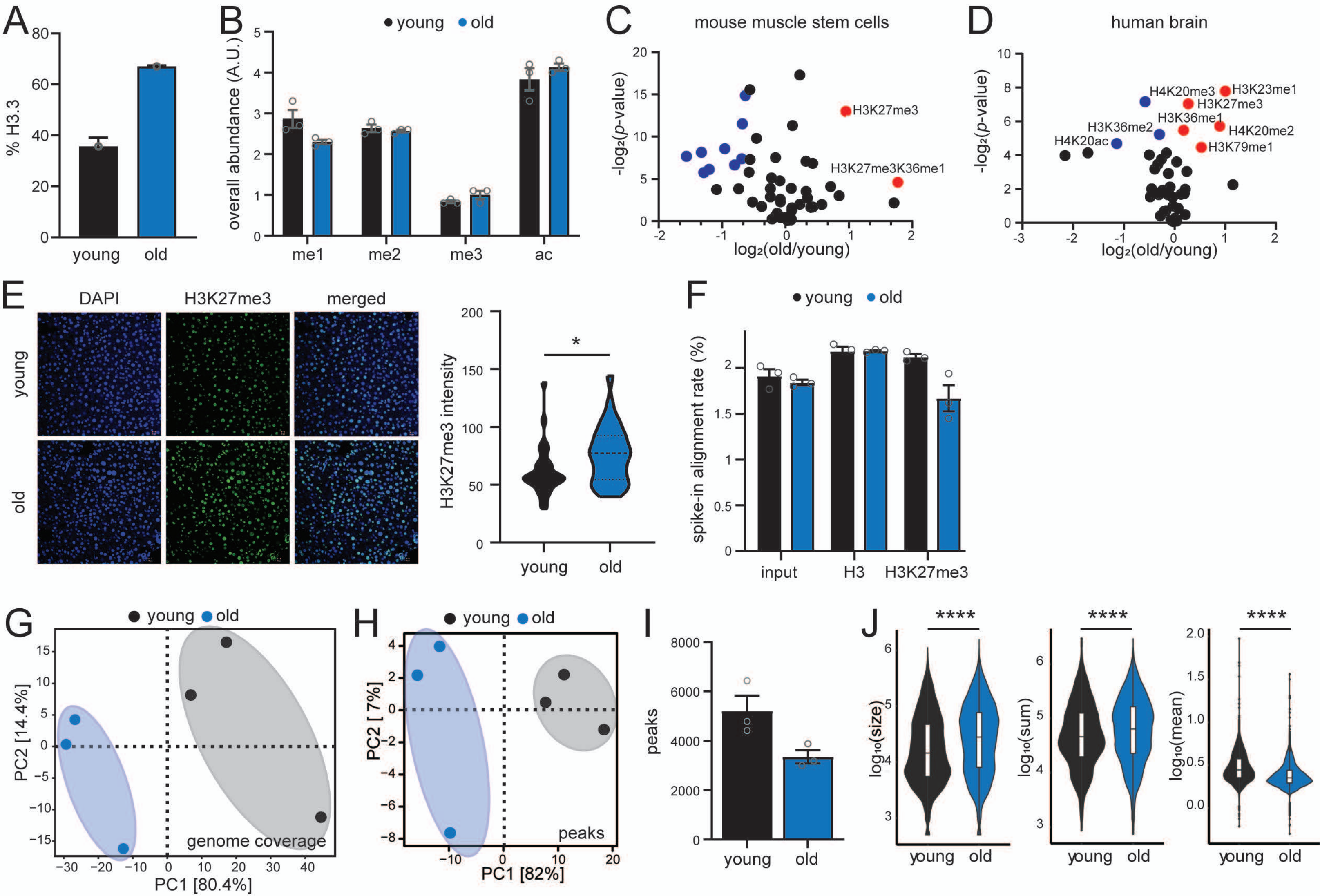


*Supplementary Figure 1*

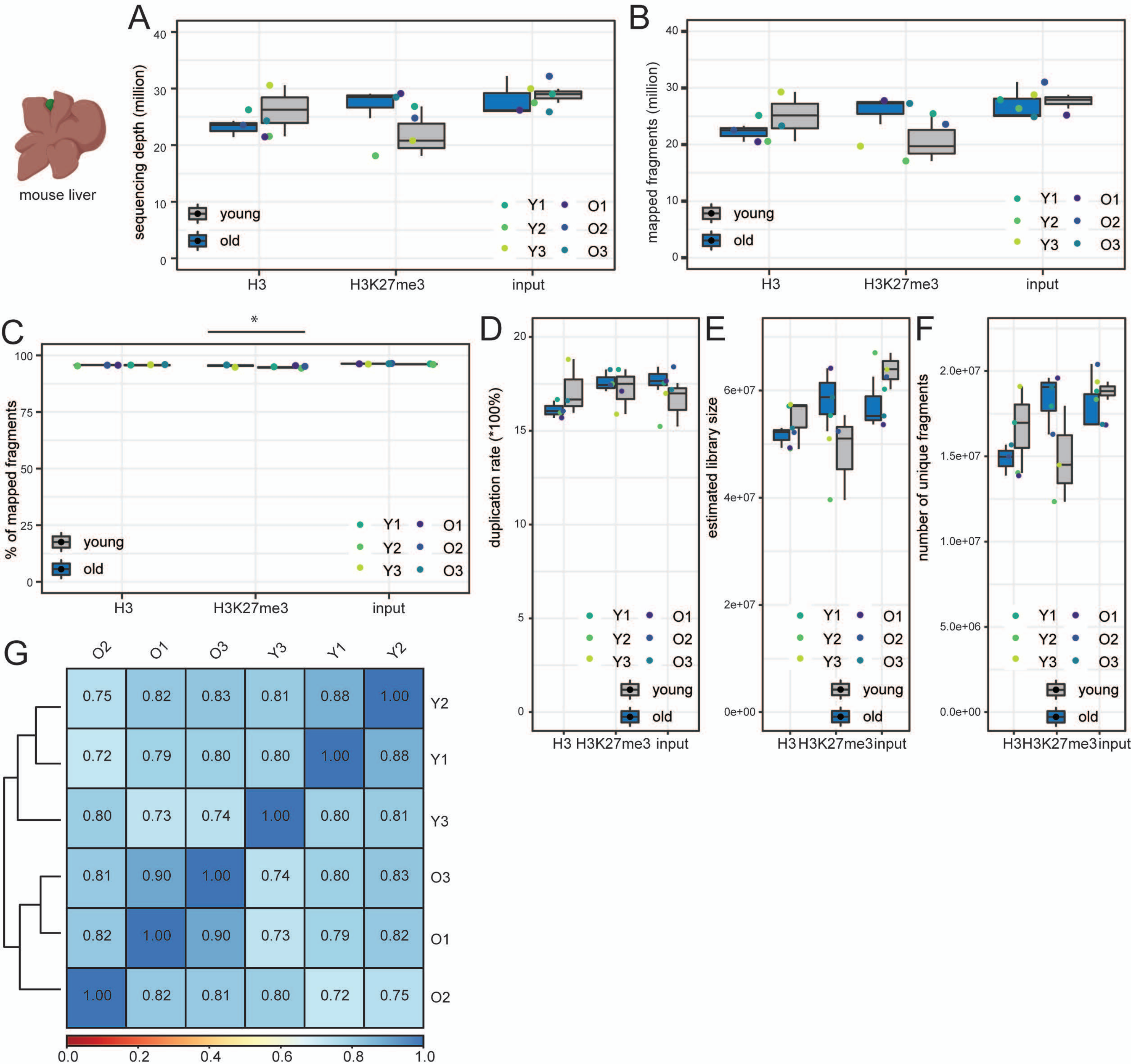


*Supplementary Figure 2*

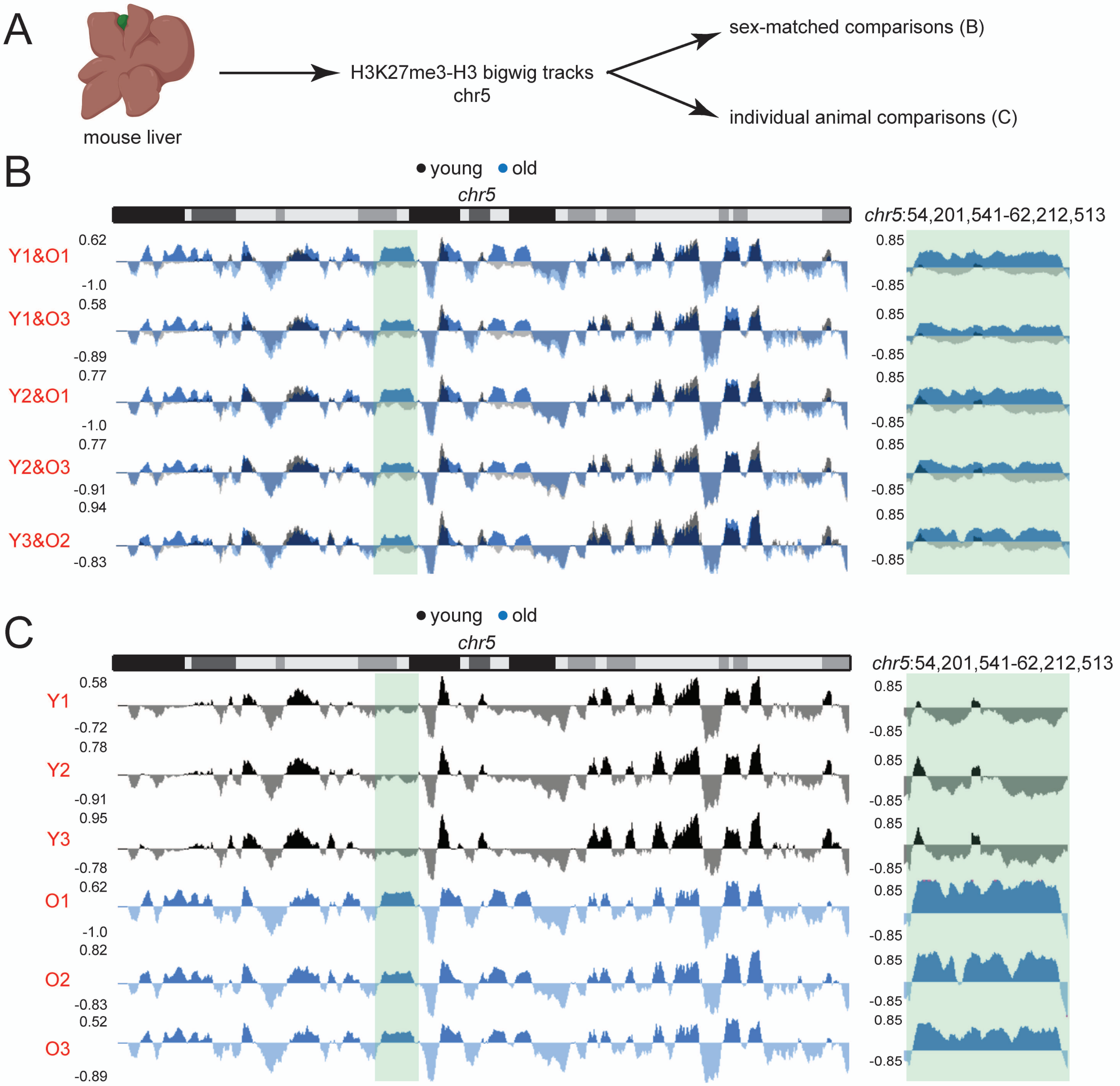


*Supplementary Figure 3*

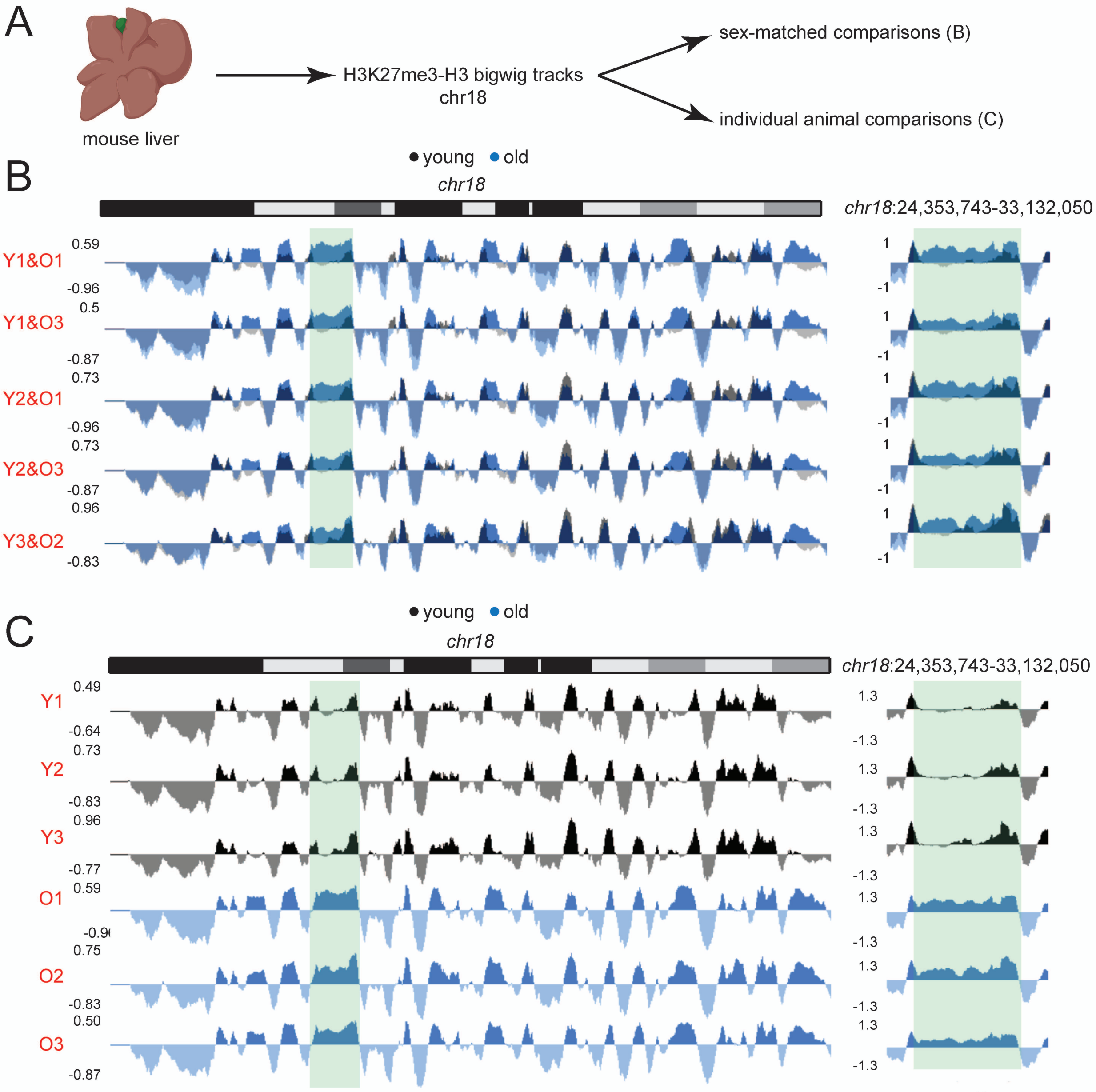


*Supplementary Figure 4*

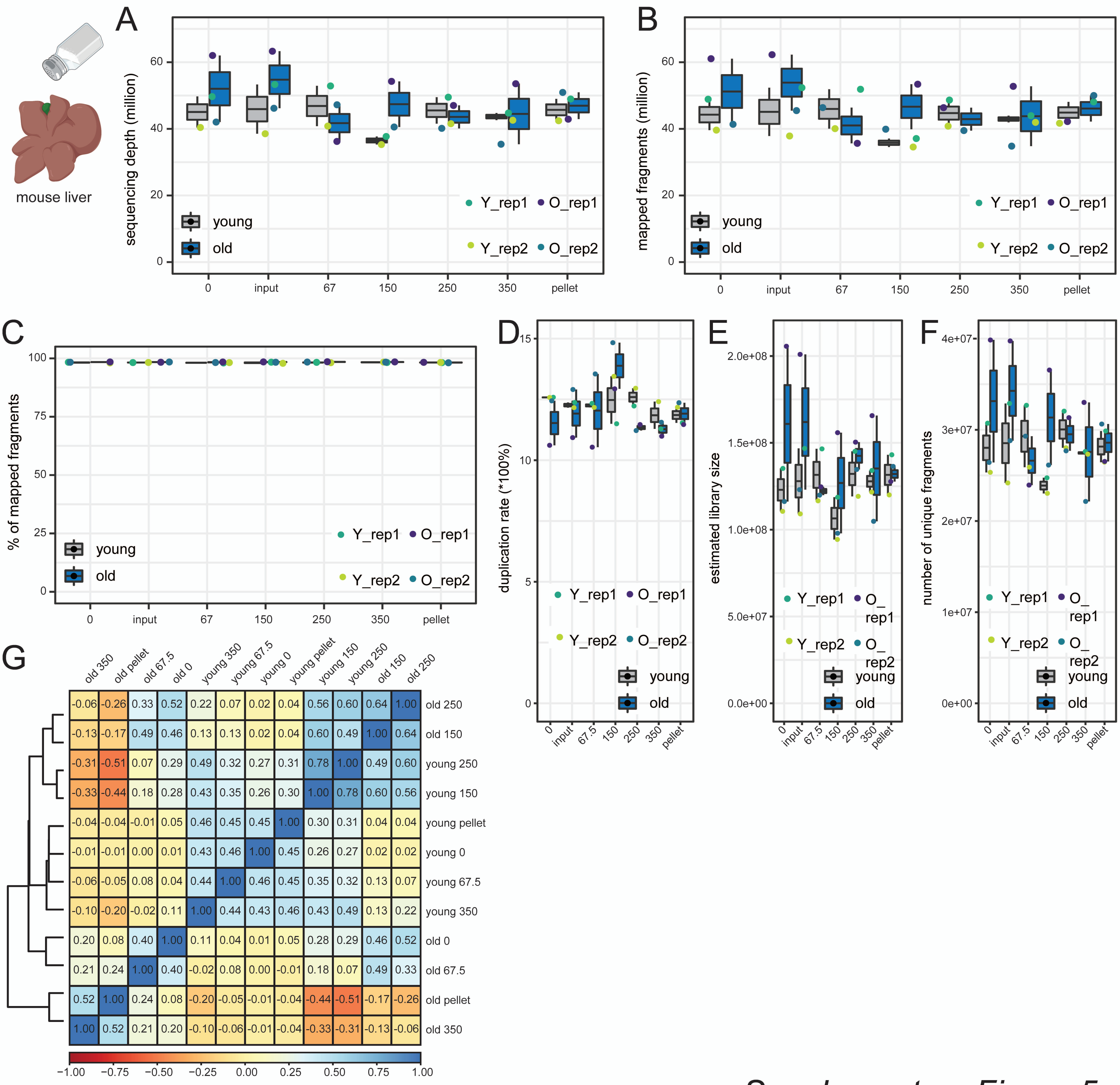


*Supplementary Figure 5*

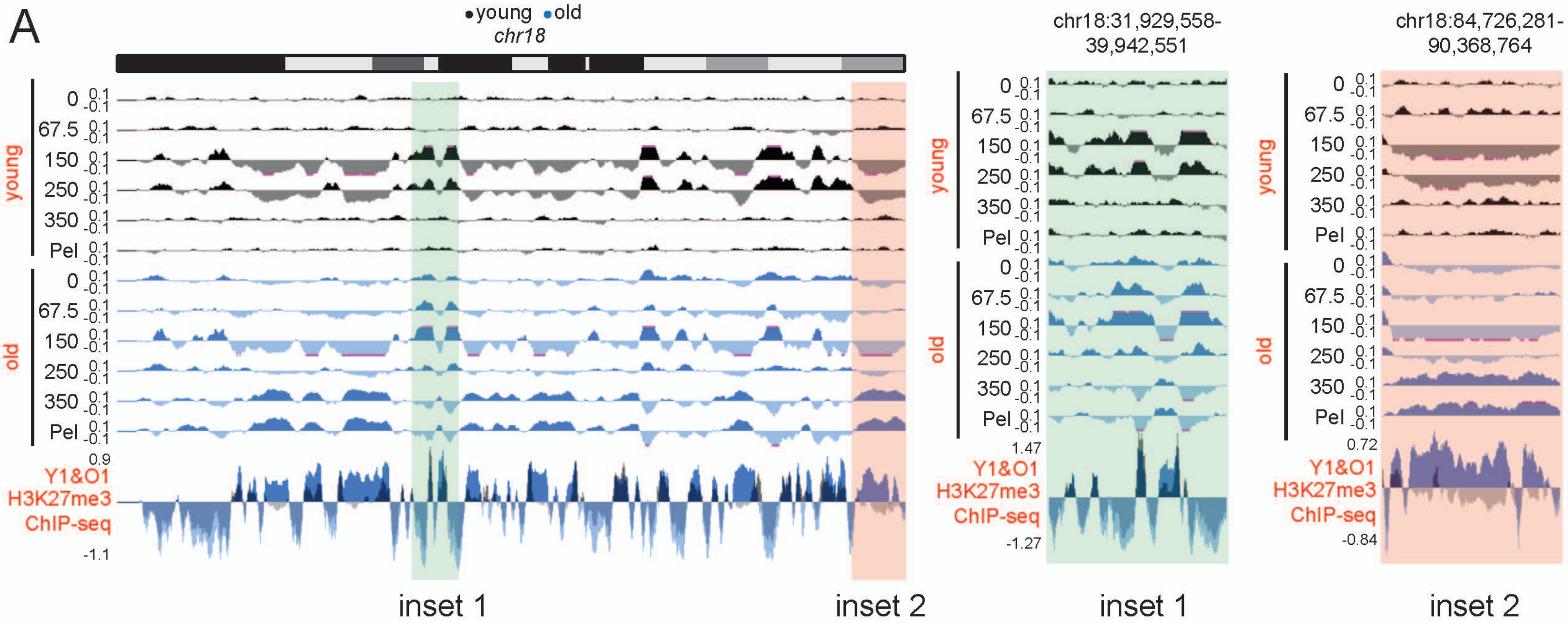


*Supplementary Figure 6*

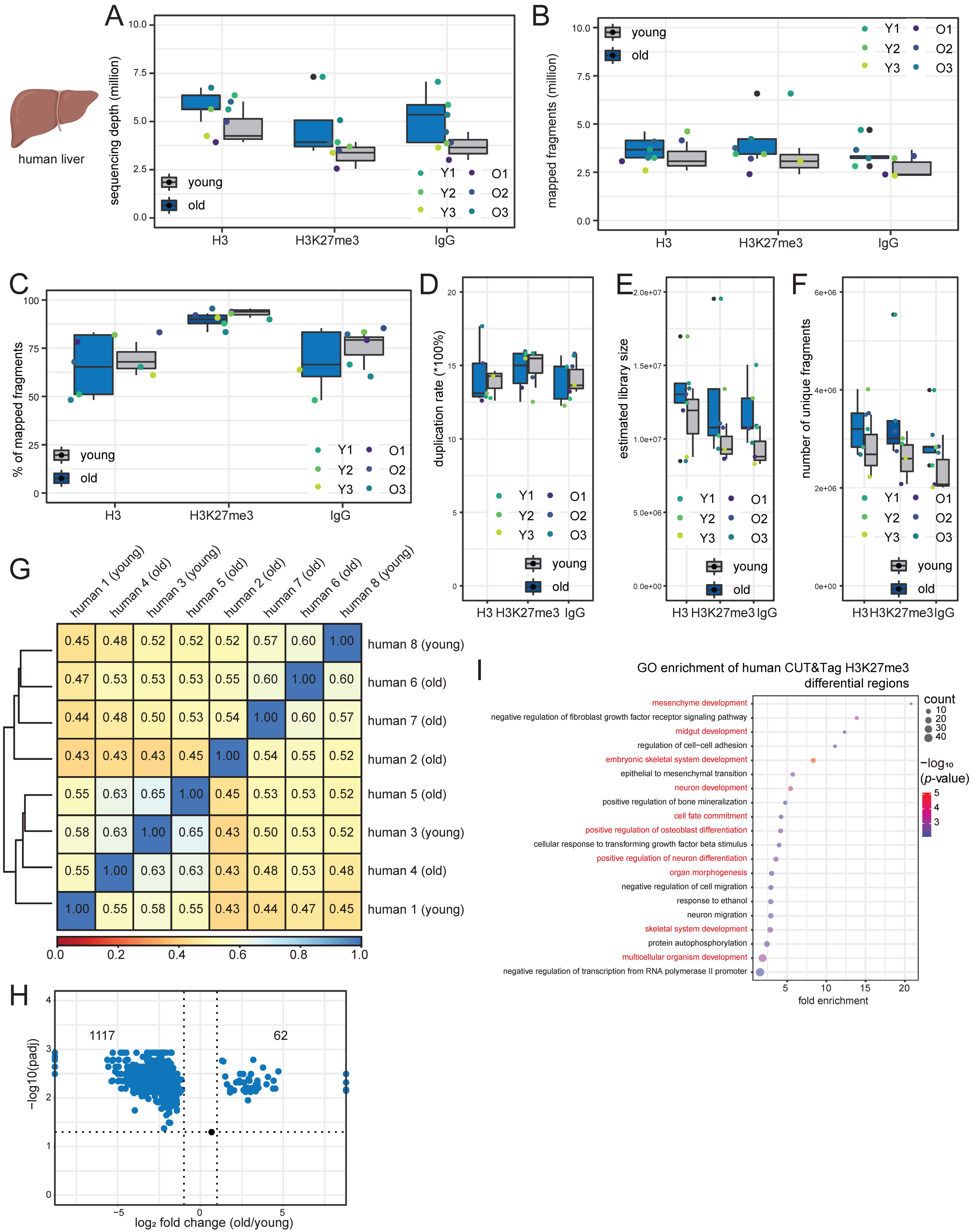


*Supplementary Figure 7*

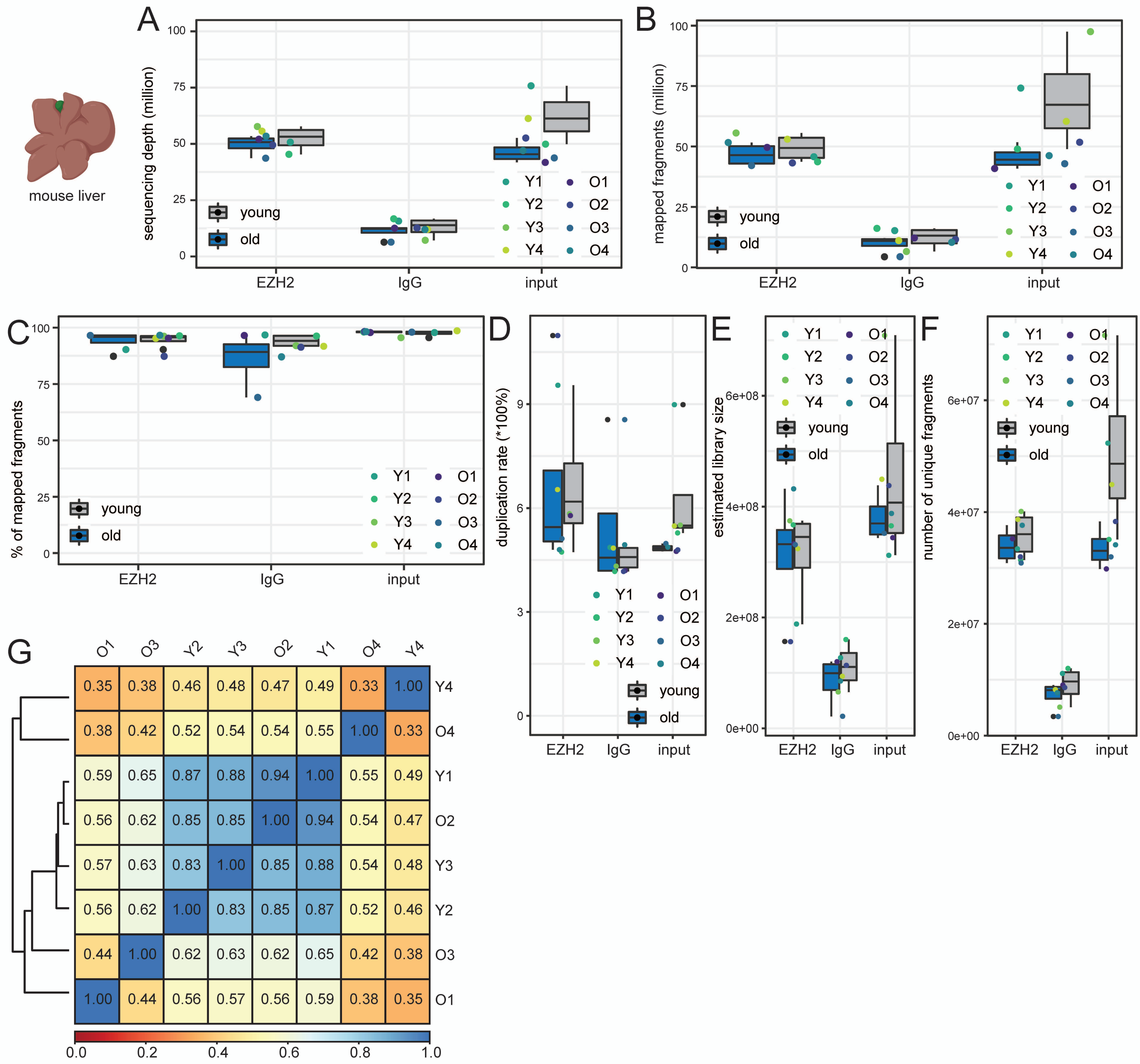


*Supplementary Figure 8*

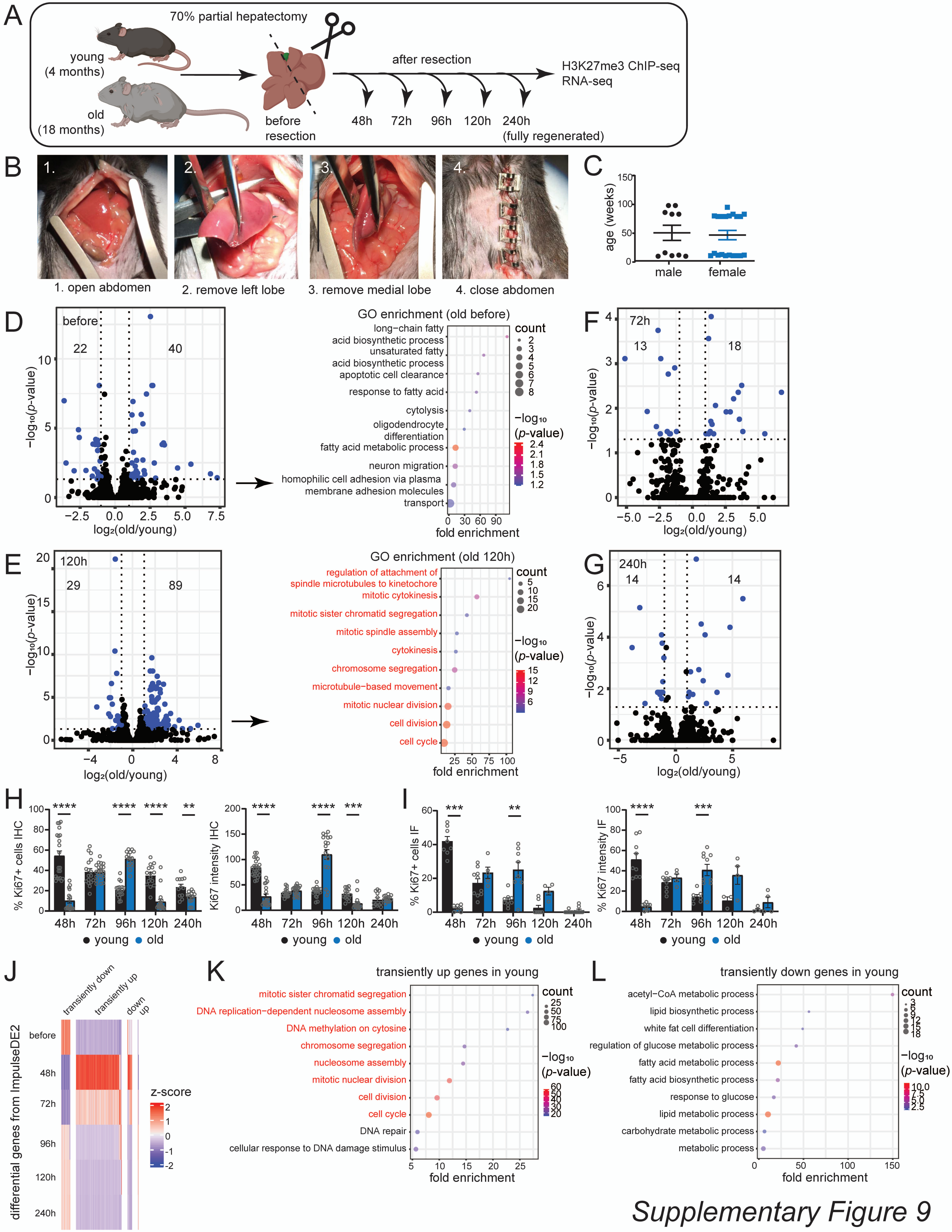


*Supplementary Figure 9*

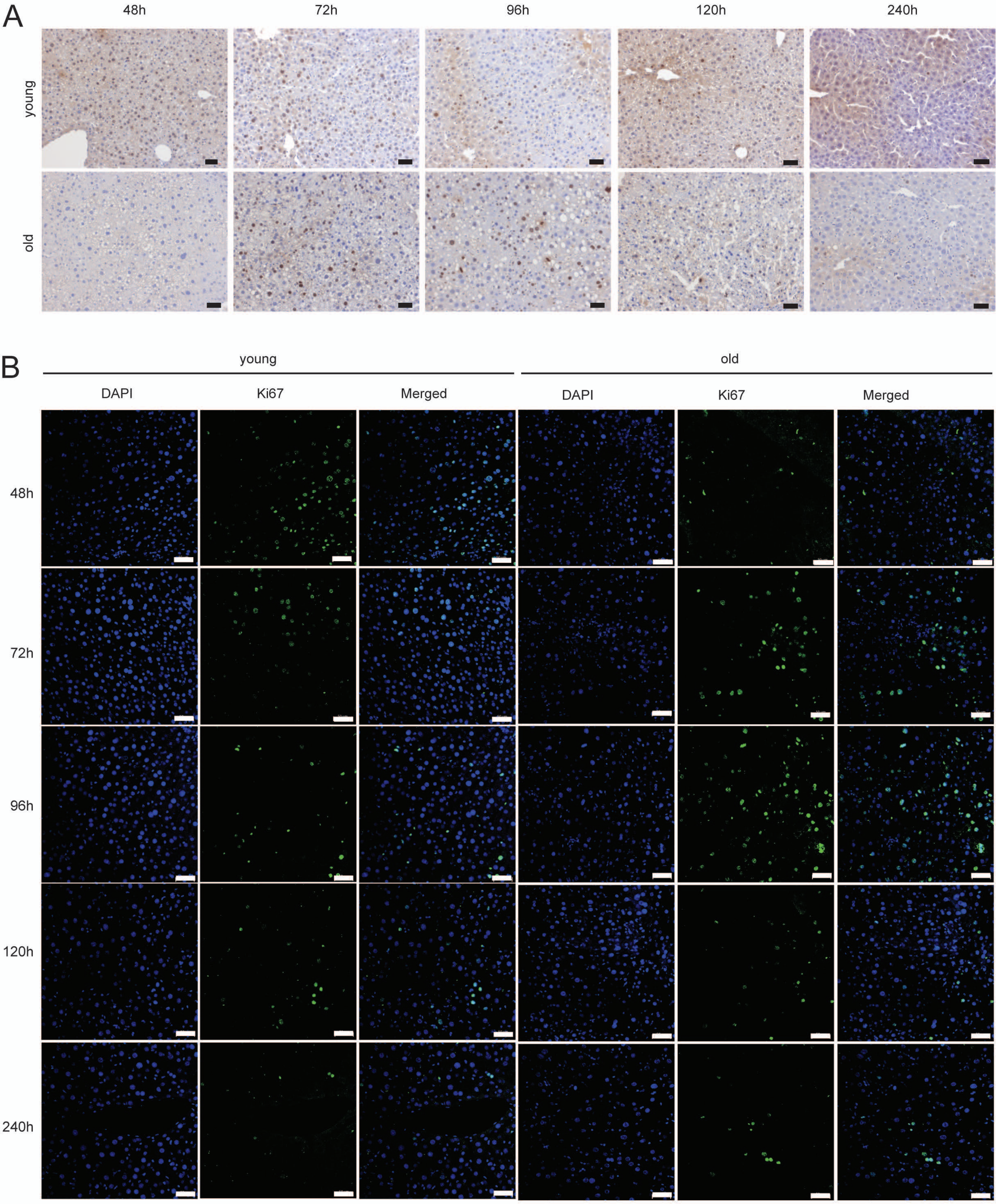


*Supplementary Figure 10*

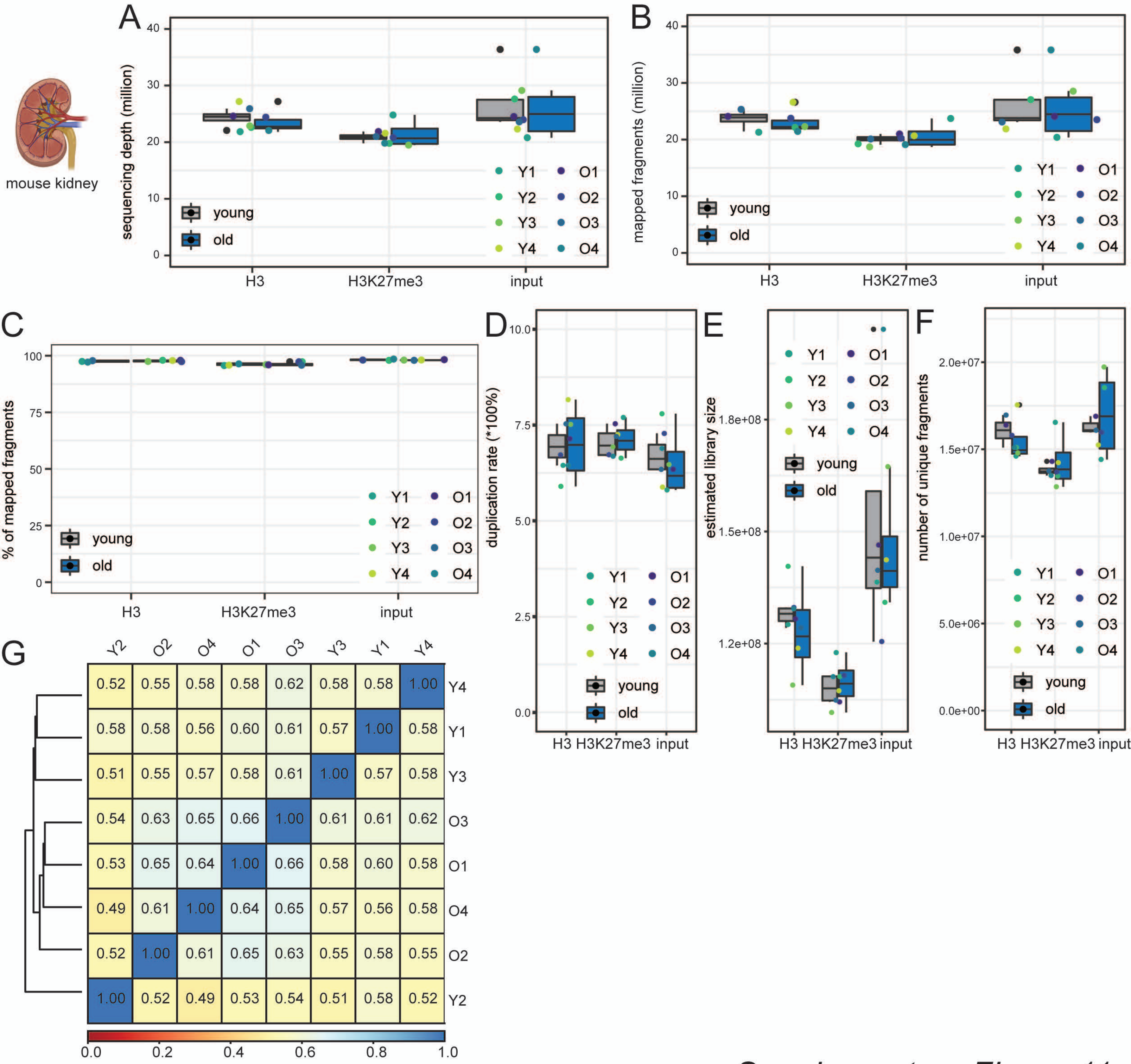


*Supplementary Figure 11*

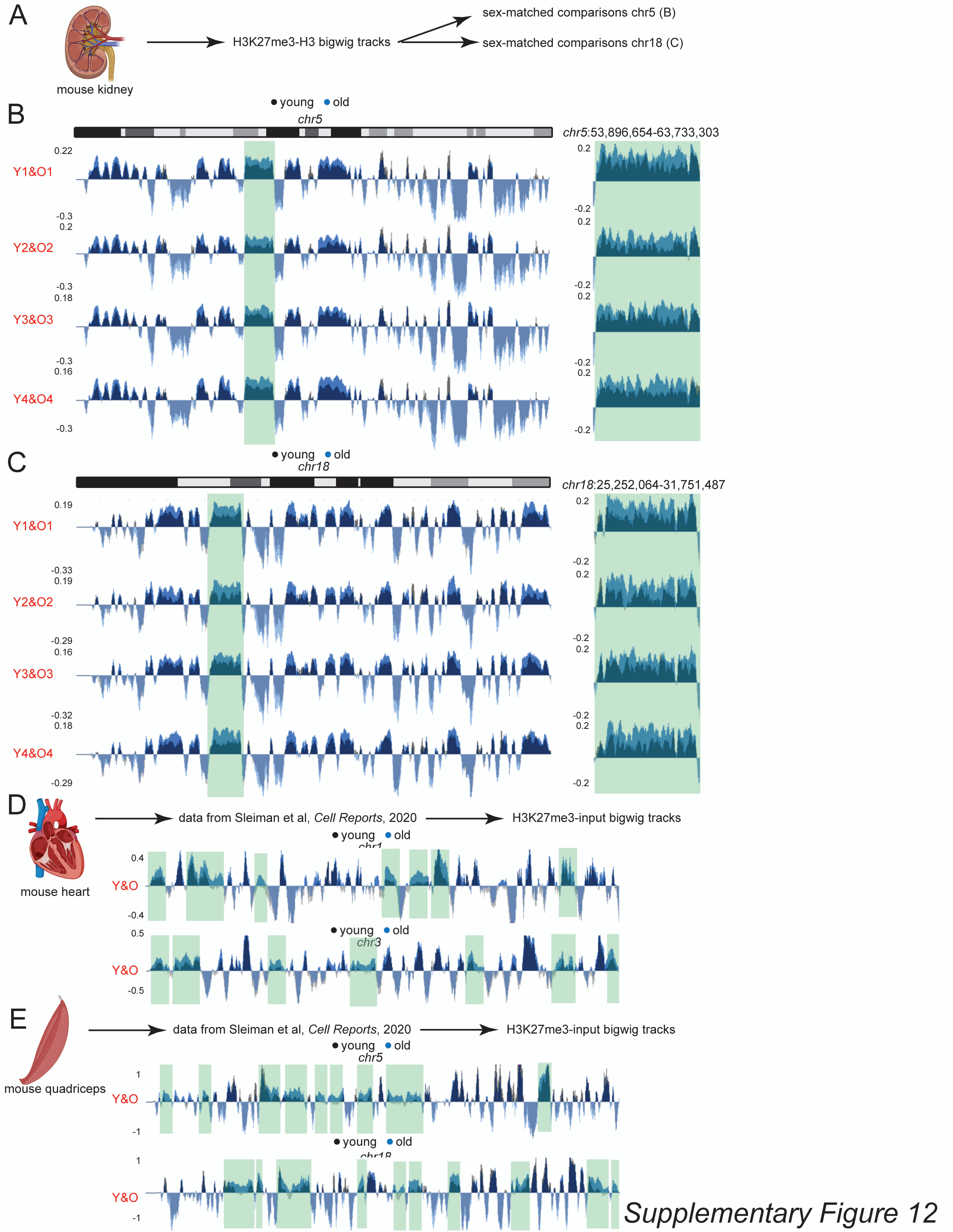


Supplementary Figure 12
